# Reduced Type-A Carbohydrate-Binding Module Interactions to Cellulose Leads to Improved Endocellulase Activity

**DOI:** 10.1101/2020.07.02.183293

**Authors:** Bhargava Nemmaru, Nicholas Ramirez, Cindy J. Farino, John M. Yarbrough, Nicholas Kravchenko, Shishir P.S. Chundawat

## Abstract

Dissociation of non-productively bound cellulolytic enzymes from cellulose is hypothesized to be a key rate-limiting factor impeding cost-effective biomass conversion to fermentable sugars. However, the role of carbohydrate-binding modules (CBMs) in enabling non-productive enzyme binding is not well understood. Here, we examine the subtle interplay of CBM binding and cellulose hydrolysis activity for three model Type-A CBMs (families 1, 3a, and 64) tethered to a multifunctional endoglucanase (CelE) on two distinct cellulose allomorphs (i.e., cellulose I and III). We generated a small-library of mutant CBMs with varying cellulose affinity, as determined by equilibrium binding assays, followed by monitoring cellulose hydrolysis activity of CelE-CBM fusion constructs. Finally, kinetic binding assays using quartz crystal microbalance with dissipation (QCM-D) were employed to measure CBM adsorption and desorption rate constants *K*_*on*_ and *K*_*off*_, respectively, towards nanocrystalline cellulose derived from both allomorphs. Overall, our results indicate that reduced CBM equilibrium binding affinity towards cellulose I alone, resulting from increased desorption rates (*K*_*off*_) and reduced effective adsorption rates (*nK*_*on*_), is correlated to overall improved endocellulase activity. Future studies could employ similar approaches to unravel the role of CBMs in non-productive enzyme binding and develop improved cellulolytic enzymes for industrial applications.

## Introduction

Microbial cellulases are broadly classified into processive exocellulases (which cleave glycosidic bonds processively from cellulose chain ends), non-processive endocellulases (which cleave glycosidic bonds within a cellulose chain), and processive endocellulases (which cleave glycosidic bonds processively upon initiating from within a cellulose chain) which work synergistically to break down cellulose (Jalak et al., 2012). t Cellulases often possess multi-modular architecture comprised of a catalytic domain (CD), which is responsible for glycosidic bond cleavage and a carbohydrate-binding module (CBM) (Talamantes et al., 2016). CBMs potentiate the activity of cognate catalytic domain against insoluble cellulose by binding to the hydrophobic face of cellulose crystals via CBM planar binding motif aromatic residues (Boraston et al., 2004), thereby increasing the local concentration of bound CD (Lehtiö et al., 2003). Previous work on processive exocellulases (like Cel7A from *Trichoderma reesei*) studied the impact of CBMs on enzyme adsorption to cellulose (Carrard et al., 2000; Fox et al., 2013; McLean et al., 2002), threading of cellulose chain through active site tunnel (Kont et al., 2016), and processive motility (Beckham et al., 2010; Brady et al., 2015). In addition, extensive research has been conducted on the reaction mechanism of processive exocellulases like Cel7A using techniques such as molecular simulations (Beckham et al., 2014; Knott et al., 2014; Vermaas et al., 2019), bulk biochemical assays (Cruys-Bagger et al., 2012; Kari et al., 2014; Kurašin and Väljamäe, 2011), single-molecule cellulase motility assays (Brady et al., 2015; Mudinoor et al., 2020; Shibafuji et al., 2014), and kinetic modeling (Levine et al., 2010; Shang et al., 2013). However, there is currently limited understanding of the elementary steps of non-processive endocellulase (used interchangeably with endocellulases hereon) action on cellulose and the impact of CBMs on overall hydrolytic activity of appended endocellulase catalytic domains.

Endocellulase action on cellulose can be explained using a simplistic model, as shown in **Figure 1**, that incorporates the following elementary steps: (i) enzyme adsorption via CBM, (ii) lateral diffusion and CD binding, (iii) complexation and hydrolysis, (iv) CD unbinding, (v) desorption via CBM. This model is based on modification of an oft-used model for exocellulases with the omission of the processive hydrolysis step (Fox et al., 2012; Mudinoor et al., 2020). This model formulation presents a unique advantage by deconvoluting the CBM adsorption/desorption steps from CD binding/unbinding and catalytic action step (Levine et al., 2010). On the subject of rate-limiting step, it was suggested that the interaction of endocellulases with so-called ‘obstacles’ on cellulose surface leads to hydrolytic rate slow-down as the reaction proceeds from burst phase to pseudo-steady state (Murphy et al., 2012). It was also suggested that this transient inactivation of endocellulases at ‘obstacles’ arises due to CBMs (Maurer et al., 2012). A recent study also showed that the predominant bound state for processive endoglucanase Cel9A was one where CBM was bound while CD was unoccupied, although these findings may not translate to other families of processive or non-processive endocellulases (Kostylev et al., 2012). This phenomenon was described in qualitative kinetic models as ‘non-productive binding’ and categorized as off-pathway (mediated by CBM) versus on-pathway (mediated by CD) non-productive binding (Gao et al., 2013b). Off-pathway non-productive binding for endocellulases collectively refers to those states where the CBM is bound to cellulose via planar aromatic residues while the CD is not catalytically engaged with substrate (see **Figure 1**). Although literature reports indicate that the attachment of CBMs to endocellulases leads to improve hydrolytic activity towards cellulose (Pan et al., 2016; Reyes-Ortiz et al., 2013), we hypothesized that tweaking the binding affinity of CBM through mutations can further improve endocellulase activity via reduced off-pathway non-productive binding. In addition, although non-productive binding states cannot be directly characterized using simple biochemical methods, measurement of CBM adsorption and desorption constants (*K*_*on*_ and *K*_*off*_ respectively) can lead to indirect insights into this phenomenon.

**Figure 1:**
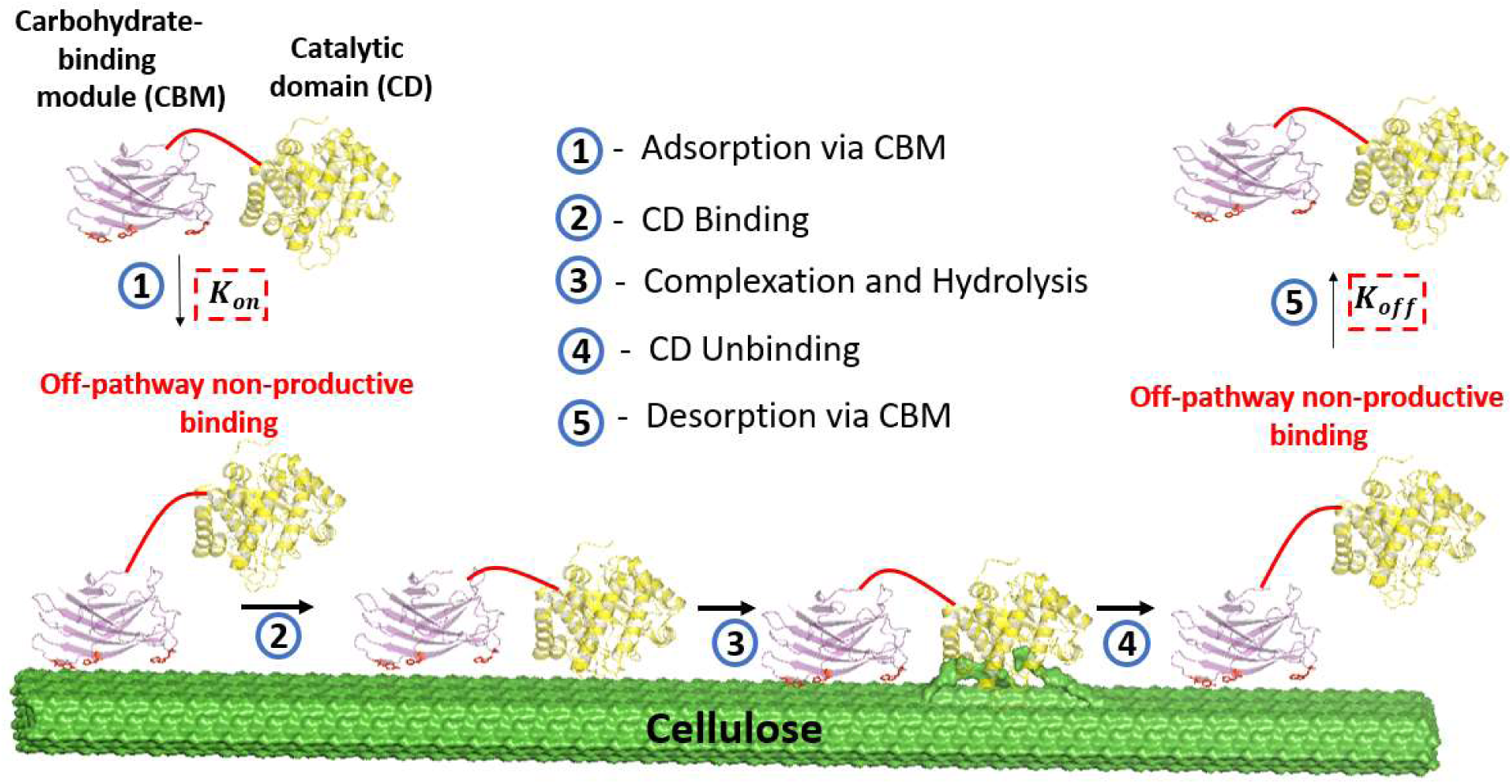
Proposed model schematic outlining steps involved in cellulose hydrolysis by multidomain endocellulases (e.g., CelE-CBM3a). (1) The first step involves enzyme adsorption mediated by the carbohydrate binding module (CBM), which can sometimes be referred to as off-pathway non-productive binding since the catalytic domain (CD) is often not engaged productively with the substrate. (2) The second step is lateral diffusion via CBM and binding of CD to cellulose substrate. (3) This is followed by complexation of a single cellulose chain to the CD active site and hydrolysis of the glycosidic bond. (4) The CD then unbinds from the cellulose surface and lateral diffusion continues until the CD binds, as shown by reverse arrow. (5) Contrarily, the enzyme can also desorb from cellulose surface into bulk solution. This model rests on the assumption that the CD adsorption/desorption kinetics from bulk solution to cellulose surface are slower than that of the CBM, which is supported by Kostylev et al. The kinetic parameters *K*_*on*_ and *K*_*off*_ outlined in this schematic refer to the adsorption and desorption constants for CBM, respectively. Published crystal structures of cellulose I fibril, CelE (PDB ID: 4IM4), and CBM3a (PDB ID: 1nbc) were used for generating this cartoon schematic using PyMOL.

Here, we seek to obtain a more comprehensive understanding of the relationship between CBM binding versus full-length endocellulase hydrolytic activity on two distinct industrially-relevant cellulose allomorphs (i.e., cellulose I and cellulose III). Cellulose III is formed during anhydrous liquid ammonia pretreatment of native cellulose I and it shows up to 5-fold increased efficiency during enzymatic hydrolysis compared to cellulose I, making it an interesting substrate for industrial adoption (Leonardo et al., 2016). We picked a model GH5 endocellulase (CelE) that has been well studied (Liu et al., 2020b; Whitehead et al., 2017), and systematically attached several mutagenized type-A CBMs to study the relationship between CBM equilibrium binding affinity and full-length endocellulase hydrolytic activity toward cellulose I and cellulose III. Finally, we measured the adsorption and desorption rate constants (*K*_*on*_ and *K*_*off*_ respectively) for CBM binding towards both cellulose allomorphs.

## Materials and Methods

See Supporting Information or SI appendix (Supplementary Text) for all materials and methods relevant to this study.

## Results and Discussion

### Native type-A CBMs show reduced binding but promote increased endocellulase activity towards cellulose III

CelE (*Ruminoclostridium thermocellum*) is a multifunctional GH5 catalytic domain with potential applications in biofuel industry due to its broad specificity toward cellulose, xylan, and mannan (Glasgow et al., 2019; Walker et al., 2017). Recently, fusing CBMs from different families to CelE improved its activity toward various pure polysaccharide substrates and pretreated lignocellulosic biomass (Walker et al., 2015). A recent study also showed that attachment of CBMs to multifunctional cellulase catalytic domains can lead to increased initial binding rates and improved biomass conversion (Brunecky et al., 2020). Here, we fused model CBMs from representative Type-A CBM families 1 (*Trichoderma reesei*) (Guo and Catchmark, 2013), 3a (*Clostridium thermocellum*) (Lehtiö et al., 2003), and 64 (*Spirochaeta thermophila*) (Pires et al., 2017; Schiefner et al., 2016) to N-terminal CelE as shown in **Figure 2A**. Gene sequences for CBM1, CBM3a, and CBM64 are provided in **SI Appendix Table T1**. Briefly, CBM1 and CBM64 genes were synthesized and sub-cloned into pEC-GFP-CBM3a and pEC-CelE-CBM3a vectors (see plasmid map in **SI Appendix Figure S1** and primers used for sub-cloning in **SI Appendix Table T2**) kindly provided by Dr. Brian Fox. The resulting fusion constructs were expressed in *E. coli* and purified to electrophoretic homogeneity. Furthermore, we studied binding and activity of GFP-CBM (GFP stands for green fluorescent protein) and CelE-CBM fusion constructs, respectively, toward Avicel based cellulose I and cellulose III (see **Figure 2B**). Preparation of Avicel cellulose III has been described in previous studies from our group (Liu et al., 2020a; Sousa et al., 2019) and the specific conditions used here for pretreatment are outlined in materials and methods section.

**Figure 2:**
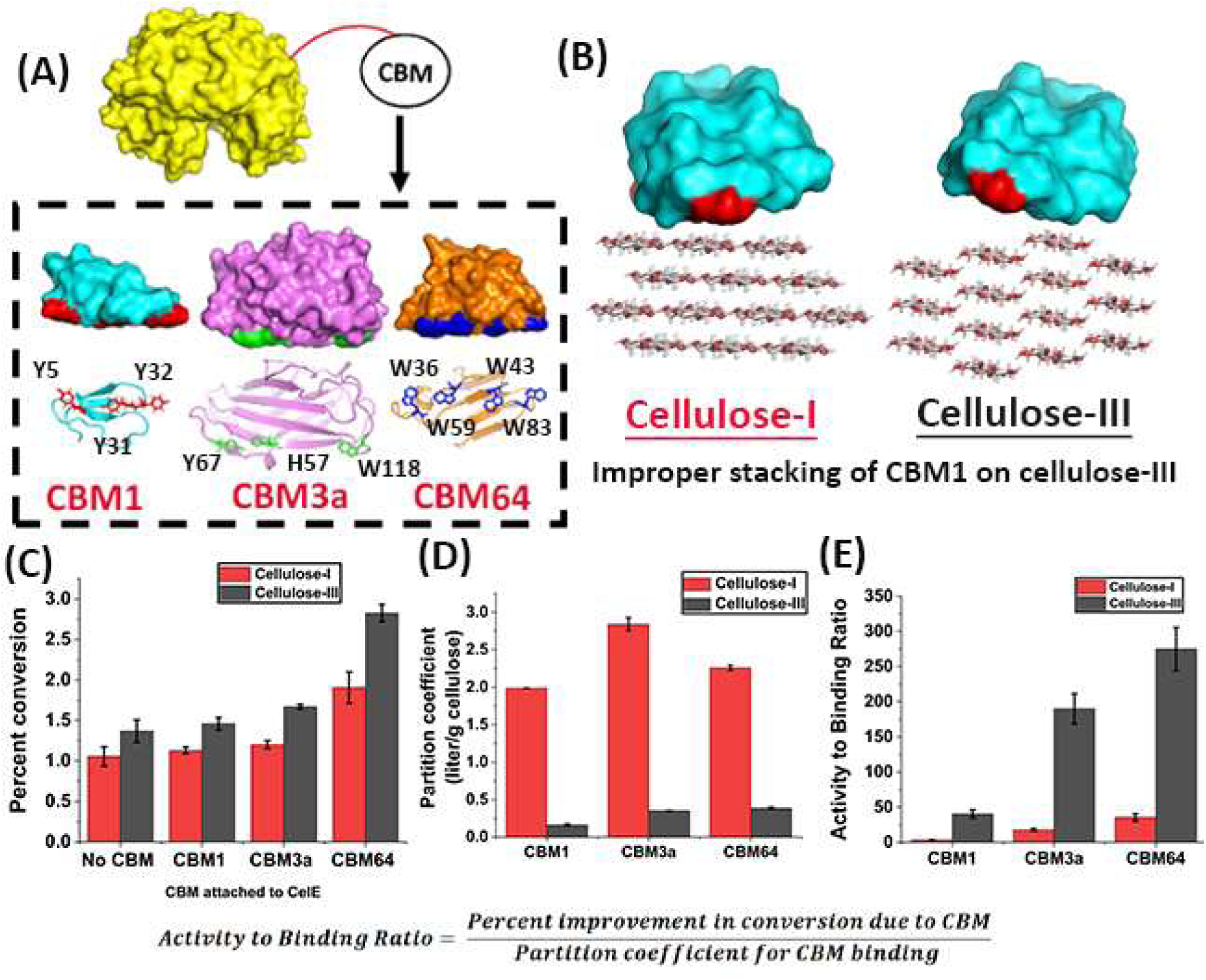
(A) Schematic of CelE-CBM fusion constructs based on crystal structures of CelE and CBMs from families 1 (*T. reesei*), 3a (*C. thermocellum*), and 64 (*S. thermophila*) generated using PyMOL. CelE is replaced by GFP to obtain GFP-CBM fusion constructs for conducting binding studies. Two perspective views of CBMs are shown here: (I) side view in surface representation (planar aromatic residues highlighted at bottom), and (II) bottom view in cartoon representation with planar aromatic residues highlighted as sticks. (B) CBM1 (*T. reesei*) shows improper stacking interactions on cellulose-III and steric clashes due to stepped cellulose-III crystal structure unlike cellulose-I. (C) Hydrolytic activity of CelE-CBM fusions and CelE control alone (no CBM) on cellulose-I (red) and cellulose-III (black) represented in terms of percent conversion of substrate to soluble sugars after 24 hours. Error bars represent standard deviation from mean based on five replicates. (D) Partition coefficient (in liter/gram cellulose) for binding of GFP-CBMs to cellulose-I (red) and cellulose-III (black). Error bars represent standard deviations from mean based on four replicates. (E) Activity to Binding Ratio was calculated based on results shown in (C) and (D) based on the formula displayed at the bottom of this figure.

CelE catalytic domain and all the CelE-CBM fusions always showed improved activity on cellulose III when compared to cellulose I as shown in **Figure 2C**. The activity on cellulose III is ∼1.3 to 1.5-fold higher with the greatest improvement observed for CelE-CBM64. These results align with recent observations for similar GH5 enzymes such as Cel5A from *Thermobifida fusca* (Liu et al., 2020b) and for other endoglucanases in general (Chundawat et al., 2011). Fluorescence based pull-down binding assays were then conducted using multiple GFP-CBM protein concentrations (ranging between 0 – 200 μg/ml) to obtain the binding partition coefficients to cellulose I and III (see **SI Appendix Figure S2** for partition coefficient raw data). These binding assay results are summarized in **Figure 2D** and indicate that native CBMs show ∼5 to 12-fold reduction in binding partition coefficient towards cellulose III. This trend is in alignment with our recent study which shows that Type-A CBMs, such as CBM1 (Chundawat et al., 2020), experience major steric clashes with the non-native surface of cellulose III due to the uneven topology of hydrophobic face as shown in **Figure 2B** that impairs CBM binding. Furthermore, CBM binding reversibility is often overlooked in the literature and this has led to contradictory results being reported (Jervis et al., 1997; Lim et al., 2014; Linder and Teeri, 1996). Here, we showed that binding of CBM families 1, 3a, and 64 to cellulose I and cellulose III is mostly reversible, ruling out the possibility of protein structural deformation on cellulose surface leading to the trends reported above (see **SI Appendix Figure S3**). However, in the case of CBM64, there was slight binding irreversibility observed towards cellulose I. To further demonstrate the relationship between CBM binding and CelE-CBM activity, we combined the data in **Figures 2C and 2D** in terms of relative activity to binding ratio which indicates the percent improvement in activity due to initial CelE catalytic activity normalized to CBM binding as shown in **Figure 2E**.

Previous studies on correlation of cellulase binding and hydrolytic activity relied either on simultaneous measurement of binding and activity (Gao et al., 2013b; Igarashi et al., 2007) or enzyme activity inhibition (by thermal denaturation or chemical inhibition) (Gao et al., 2011; Jung et al., 2003). However, these approaches do not enable clear demarcation of CBM-driven binding interactions versus CD-driven binding interactions which can be achieved by studying CBM binding alone. Overall, these results indicate that attaching a Type-A CBM to CelE leads to greater hydrolytic activity per unit bound CBM toward cellulose III versus cellulose I. Based on this finding, we wanted to further examine whether a reduction in CBM binding to either substrate (i.e., cellulose I or cellulose III), could lead to a similar improvement in hydrolytic activity as seen for the native CelE-CBM fusion constructs. To test this hypothesis, we mutagenized the CBM planar binding motif aromatic residues to alanine.

### Planar binding motif aromatic residue mutations lead to a reduction in binding affinity towards both cellulose I and cellulose III

CBM1 was excluded from this mutagenesis study because previous studies have predicted structural deformation of this rather small protein (∼36 kDa) when key residues such as Y5 were mutated to alanine (Pettersson et al., 1995). In addition, the impact of mutations is likely to be highly dependent on glycosylation due to fungal origin of CBM1 (Guan et al., 2015; Taylor et al., 2012). Site-directed mutagenesis and protein production/purification were performed as described in the materials and methods section, to generate GFP-tagged CBM3a and CBM64 mutants. Fluorescence based full-scale GFP-CBM/cellulose binding assays were conducted and the resulting data was fit to Langmuir isotherm one-site model to obtain the total number of available cellulose surface binding sites (*N*_*max*_) and dissociation constant (*K*_*d*_) (see raw data and model fits reported in **SI Appendix Figure S4-S5**). Regardless of the substrate, the mutants always showed a reduction in binding affinity (i.e., inverse of dissociation constant *K*_*d*_) compared to the respective wild-type proteins (used interchangeably with WT hereon) (see **Table 1**). For cellulose I, CBM3a mutants showed a reduction in binding affinity ranging from ∼8 to 33-fold, with Y67A mutant showing the least affinity. CBM64 mutants showed a reduction in binding affinity ranging from ∼5 to 16-fold, with W36A ranking the least. For cellulose III, CBM3a mutants showed ∼3 to 8-fold reduction with the rank order being Y67A > H57A ∼ W118A; whereas CBM64 mutants showed ∼3 to 28-fold reduction with W36A showing the least affinity. The number of binding sites (*N*_*max*_) for mutants showed a slight improvement or remained the same for most mutants on cellulose I except for CBM3a-W118A and CBM64-W36A. In addition, for any given mutant, the number of binding sites on cellulose III was always lesser compared to cellulose I. The mutations also reduced the estimated apparent partition coefficient 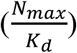 on both substrates as reported in **Table 1**.

**Table 1:**
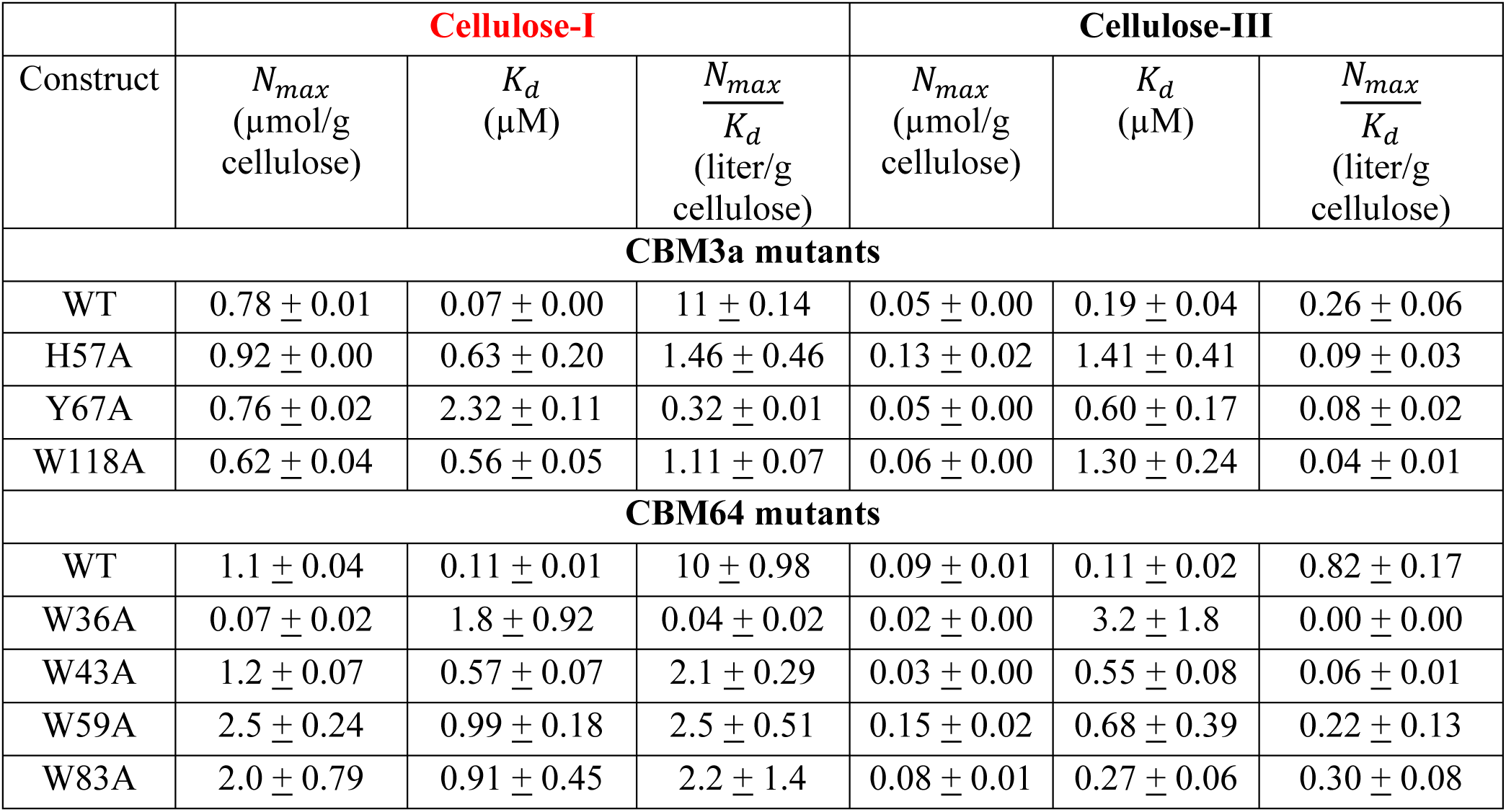
Binding parameters *N*_*max*_, *K*_*d*_ for GFP-CBM3a and GFP-CBM64 mutants obtained from Langmuir one-site model fits to full-scale binding assay data. Errors reported here are standard errors from the mean obtained from fitting analysis. The experiments were performed using six replicates for each protein concentration as reported in **SI Appendix Figures S4-S5**. Here, WT-wild-type for each respective CBM family.

Here, we used fluorescent protein tagging to study binding of CBMs to cellulose as also previously demonstrated (Hong et al., 2007; Novy et al., 2019). Researchers have previously shown that mutation of type-A CBM planar aromatic residues can reduce binding to cellulose I which aligns with our results (McLean et al., 2000; Nagy et al., 1998; Pettersson et al., 1995; Reinikainen et al., 1995). To the best of our understanding, this is the first study to have demonstrated the role of these aromatic residues in CBM binding to cellulose III. Qualitatively, these mutations seem to impact binding to both cellulose I and cellulose III in a similar manner by reducing binding affinity. It is likely that the loss of hydrophobic stacking interactions between the aromatic residues and cellulose chains impacts binding to both allomorphs (Chundawat et al., 2020; Georgelis et al., 2012). To further understand the impact of these mutations on hydrolytic activity, we produced and characterized the CelE-CBM mutant constructs as described below.

### CBM planar binding motif aromatic residue mutations lead to improved endocellulase activity on cellulose I

Site-directed mutagenesis, protein expression, and purification to generate CelE-CBM3a and CelE-CBM64 mutants, were performed as described in the materials and methods section. Activity assays were then performed on the mutants and the resultant activities are reported in the form of a parameter called percent relative activity, which compares mutant activity to wild-type (WT) activity (see **Figure 3**). Surprisingly, every mutation improved the activity on cellulose I significantly ranging from ∼20 to 70% improvement for various CelE-CBM3a (**Figure 3A**) mutants and ranging from ∼6 to 80% improvement in the case of CelE-CBM64 mutants (**Figure 3B**). Towards cellulose III, in contrast, there was only a marginal improvement in the case of all CelE-CBM3a mutants (except for CBM3a-W118A mutant) and activity reduction was seen in all cases with CelE-CBM64 mutants (except for CBM64-W43A mutant). One of the flanking aromatic residues in each case, W118 (for CBM3a) and W36 (for CBM64) (**Figure 2A**) showed the least improvement in activity on cellulose I, implying the importance of these residues in aligning the CBM properly on the cellulose I surface for productive catalysis by the CD. We also tested the impact of enzyme loading and reaction time on cellulose I and cellulose III activities but did not notice a difference on relative trends (see **SI Appendix Figure S6**). Furthermore, we chose 24 hours as the reaction time for all our activity assays because longer reaction times can lead to protein denaturation and hence irreversible binding. Overall, correlating these results to the affinity of CBM mutants reported in previous section, we find that reducing binding affinity of CBMs while maintaining the number of binding sites improves catalytic activity of corresponding CelE-CBM mutants toward cellulose I but not cellulose III.

**Figure 3:**
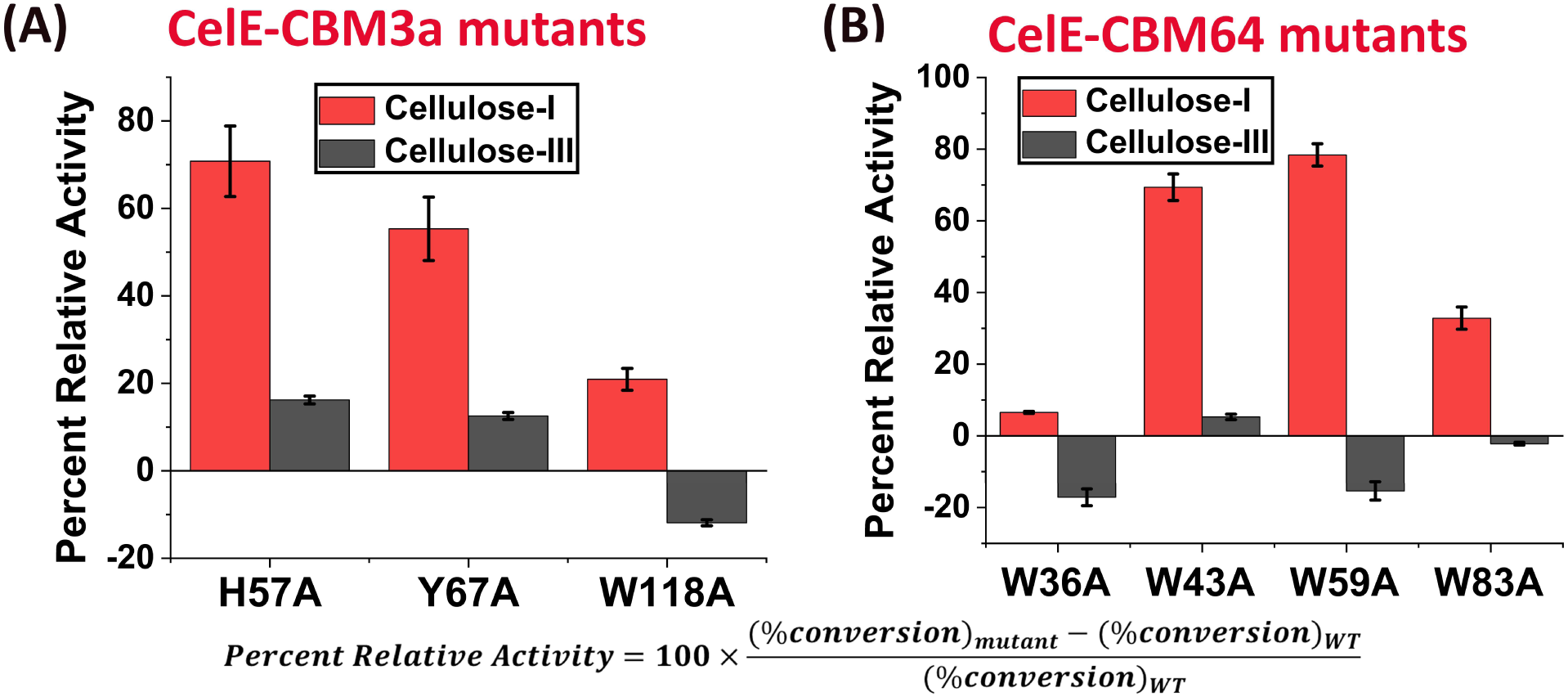
(A) Percent relative activity for CelE-CBM3a mutants calculated based on hydrolytic activity of mutants in comparison to activity of wild-type (WT) on a given substrate: cellulose-I (red) or cellulose-III (black). Percent relative activity is calculated based on the formula at the bottom of this figure. (B) Percent relative activity for CelE-CBM64 mutants towards cellulose-I (red) and cellulose-III (black). Error bars represent standard deviation from the mean based on five replicates.

Previous literature reports have shown that fusion of CBMs to endocellulases from GH5 family improves activity toward native cellulose I (Arumugam Mahadevan et al., 2008; Sajjad et al., 2012; Yoda et al., 2005). Inter-domain interactions between CBM and CD were also shown to play a critical role in the case of GH9 endoglucanases (Burstein et al., 2009), leading to a reduction in activity when an aromatic residue on CBM3c was mutated (Kim et al., 2016). It is unlikely that specific interactions exist between CelE and CBM3a/64 since they did not coevolve tethered together. Hence, we speculate that the improved activity of CelE-CBM3a and CelE-CBM64 mutants toward cellulose I arose majorly from reduced binding of CBMs to cellulose although future experiments are needed to fully validate this claim. In addition, type-A CBMs (such as CBM1/3a/64) bind ordered crystalline regions (Novy et al., 2019) whereas endocellulases like CelE target amorphous regions of cellulose (Orłowski et al., 2015). We also speculate that this disparity in substrate specificities between type-A CBM and endocellulase CD leads to off-pathway non-productive binding which can be mitigated by reduction of CBM binding affinity towards cellulose I. On the other hand, cellulose III shows a dramatic reduction in the number of binding sites for wild-type proteins when compared to cellulose I. Our results on cellulose III show that further reduction of binding sites and affinity upon CBM mutation leads to marginal or no improvement in activity, possibly because the hydrolysis of this allomorph is limited by the amount of bound enzyme under the specific conditions used in this study. As a result, protein engineering efforts for activity improvement on cellulose III need to be focused on engineering the CD or improving affinity of CBM.

### QCM-D assays indicate differences in kinetic behavior of CBM mutants towards cellulose I versus cellulose III

Quartz crystal microbalance with dissipation (QCM-D) binding assays were setup to measure CBM binding kinetics as shown in the schematic (**Figure 4A**). Briefly, the frequency data acquired during binding and unbinding of proteins was converted to number of protein molecules using the Sauerbrey equation (Brunecky et al., 2020), as shown in **Figure 4B**. The raw data for frequency and dissipation changes during CBM3a and CBM64 wild-type (WT) binding to cellulose I and cellulose III is shown in **SI Appendix Figure S7**. The unbinding regime specifically, was fit to an exponential decay equation to obtain a true desorption rate constant *K*_*off*_ (see materials and methods section for model-fitting equations). All parameters from fitting analysis aside from *K*_*off*_ have been reported in **SI Appendix Table T3**. Effective adsorption rate constant *nK*_*on*_ was measured using *K*_*off*_ from QCM-D binding assays and *N*_*max*_, *K*_*d*_ obtained from equilibrium binding assay results (see **Figure 4C** for the formula used). These kinetic parameters were specifically used since they were an integral part of the qualitative cellulase binding-activity kinetic model that we reported previously (Gao et al., 2013a). Compared to WT, CBM3a mutants showed ∼ 5 to 25-fold lower *nK*_*on*_ towards cellulose I and an increased *K*_*off*_ with a maximum ∼1.4-fold increase observed for Y67A (**Figure 4C (I, II)**). A similar trend was observed for all the CBM64 mutants towards cellulose I (**Figure 4D (I, II))** with *nK*_*on*_ lowered by ∼ 2 to 34-fold and *K*_*off*_ increased by ∼2 to 8-fold compared to the respective values for WT. W59A was the only exception amongst CBM64 mutants as this mutation led to a ∼ 1.5-fold reduction in *K*_*off*_ as opposed to an increased *K*_*off*_ observed in all other cases. In summary, planar aromatic residue mutations for both CBM3a and CBM64 led to a significant reduction in *nK*_*on*_ and a concomitant increase in *K*_*off*_ toward cellulose I in most cases, which correlated with the increased activity for corresponding CelE-CBM mutants compared to their WT (see **Figure 3**).

**Figure 4:**
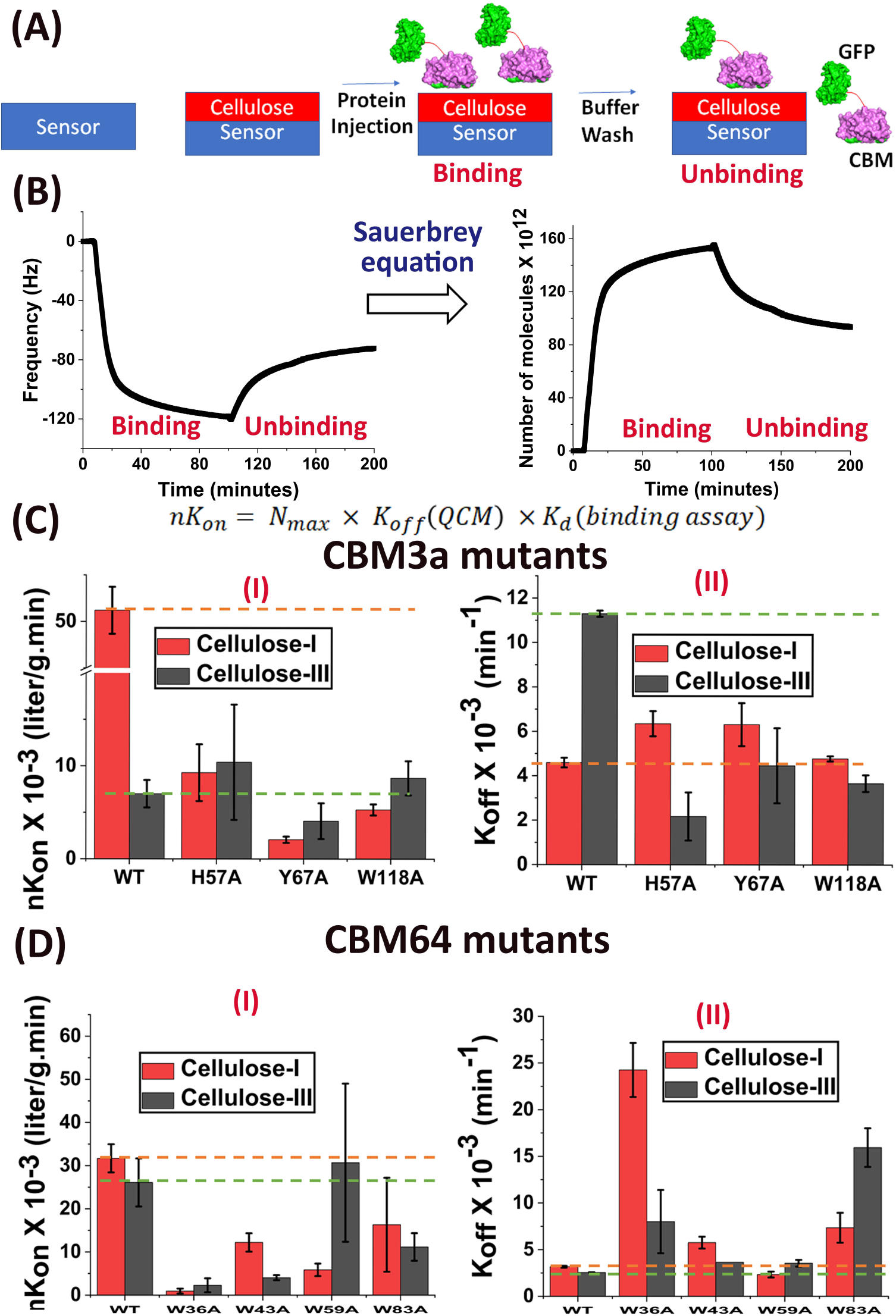
(A) Schematic for QCM-D based CBM-cellulose binding assay. (B) Frequency (Hz) vs time (minutes) data for a representative protein (GFP-CBM3a WT) was converted to number of protein molecules X 10^12^ vs time (minutes) (also called sensorgram) using Sauerbrey equation. The binding and unbinding data in plot on right was then fitted to an exponential rise and decay function, respectively, as described in detail in the **SI Appendix**. (C) (I) nK_on_ was calculated using the formula stated and nK_on_, and (II) K_off_ for CBM3a mutants toward cellulose-I (red) and cellulose-III (black) is reported here. (D) (I) nK_on_ and (II) K_off_ for CBM64 mutants toward cellulose-I (red) and cellulose-III (black) is reported here. Error bars in (C) and (D) represent standard deviations from the mean based on two replicates. Green and orange dotted lines in (C) and (D) are drawn to guide the eye towards binding parameters for wild-type (WT) control GFP-CBM towards cellulose-I and cellulose-III, respectively. In (D-II), the orange and green dotted lines are indistinguishable since both cellulose I and cellulose III have similar values. In (C) and (D), WT – wild-type for each respective CBM family.

However, the kinetic behavior of CBM3a and CBM64 mutants toward cellulose III was very different from that observed for cellulose I. CBM3a mutants showed *nK*_*on*_ similar to the wild-type protein whereas *K*_*off*_ was lowered by ∼2.5 to 5-fold. On the other hand, CBM64 mutants showed marginally lower or similar *nK*_*on*_ and increased *K*_*off*_ on cellulose III compared to the wild-type (WT). In summary, somewhat similar or increased *nK*_*on*_ coupled with reduced *K*_*off*_ correlated with increased activity of CelE-CBM3a mutants with the exception of W118A. Conversely, lower *nK*_*on*_ and somewhat increased or similar *K*_*off*_ correlated to reduced activity for CelE-CBM64 mutants. When interpreted in light of our proposed model for endocellulase action, these results suggest that off-pathway non-productive binding might be prevalent in the case of cellulose I but not cellulose III, which could be mitigated by lowering *nK*_*on*_ and increasing *K*_*off*_ for CBM binding to cellulose.

QCM-D has been employed previously to monitor enzymatic hydrolysis of cellulose films (Maurer et al., 2013; Turon et al., 2008), study binding of full-length cellulases to cellulose allomorphs (Brunecky et al., 2020; Cheng et al., 2012; Hu et al., 2018; Maurer et al., 2012), and study binding of cellulases to lignin and pretreated biomass (Haarmeyer et al., 2017; Kumagai et al., 2014; Sammond et al., 2014). Here, we also attempted to capture the viscoelastic behavior of CBM-cellulose binding process using a parameter called specific dissipation (see **SI Appendix Figure S8-S9**). Marginally increased specific dissipation was observed for all proteins tested on cellulose I vs cellulose III, except for CBM3a-H57A. Previous reports suggest that increased specific dissipation in case studies of protein adsorption could arise either from increased film hydration (Höök et al., 2001; Tammelin et al., 2015) or due to structural deformation of protein (Jordan and Fernandez, 2008). An increase in conformational entropy either due to solvent release or protein deformation was correlated to dissipation increases in the case of HIV protein-ligand binding interactions (Lee et al., 2008). It is likely that the release of interfacial water molecules upon CBM binding (Georgelis et al., 2012; Orłowski et al., 2018) leads to increased dissipation for cellulose I versus cellulose III. The hydrophobic binding face of cellulose III was previously shown to form ∼1.5-fold greater hydrogen bonds with water leading to favorable enthalpic versus entropic interactions (Chundawat et al., 2011). These results show that QCM-D can be used to infer useful information on hydration and protein deformation patterns of CBM-cellulose interactions, in addition to binding kinetics.

## Conclusions

CBMs play an important role in targeting glycoside hydrolase enzymes to plant cell wall polysaccharides such as cellulose (Fox et al., 2013; Herve et al., 2010; McLean et al., 2002). However, the importance of appended CBMs to cellulase activity has been recently brought into question by their apparent insignificance under high solids biomass loadings in biorefinery settings (Pakarinen et al., 2014; Varnai et al., 2013). In addition, CBMs have been predicted to lead to non-productive binding of cellulase enzymes to cellulose although predictions from these kinetic models were not proven experimentally (Gao et al., 2013a; Nill and Jeoh, 2020). Moreover, the impact of CBM fusion on activity of commercially relevant cellulases toward pretreated substrates is not well understood either (Kim et al., 2010). In this study, we show that reducing the strength of type-A CBM binding to native cellulose I leads to an increase in the activity of model endocellulase CelE fusion enzymes in the pseudo steady-state regime. In summary, these results provide incremental supporting evidence to our original hypothesis that off-pathway non-productive binding can be mitigated by mutagenizing CBMs, which leads to overall improved activity on cellulose I. These results also align with the recently proposed Sabatier principle, which posits that increased cellulase binding to substrate is not always optimal for catalysis (Kari et al., 2018). Cellulose III, on the other hand, showed marginal to no improvement upon reduced CBM binding indicating that the hydrolysis of this allomorph may be limited by the amount of bound endocellulase enzymes.

Although the results from this study shed light on the interplay of CBM binding affinity and binding kinetics versus hydrolytic activity, similar binding/activity studies need to be performed on a larger library of full-length enzymes to draw general conclusions for the full catalytic cycle of endocellulases. Additionally, future studies employing pre-steady state enzyme kinetic assays could be also employed to contrast rate limitations for diverse endocellulase families toward more industrially relevant substrates like cellulose III versus native cellulose I (Kari et al., 2018; Kari et al., 2020). Lastly, more detailed multiscale simulation studies on distinct cellulose surfaces will be needed to fully understand the rate-limiting steps of CBM-tethered endocellulases over the entire non-processive catalytic cycle as currently hypothesized in this study and elsewhere (Bianchetti et al., 2013; Orłowski et al., 2015).

## Acknowledgements

The authors acknowledge support from the NSF CBET awards (1604421 and 1846797), ORAU Ralph E. Powe Award, Rutgers Global Grant, Rutgers Division of Continuing Studies, Rutgers School of Engineering, and the Great Lakes Bioenergy Research Center (DOE BER Office of Science DE-FC02-07ER64494). SPSC and BN would like to particularly thank Professor Brian Fox (UW Madison) and Professor Bruce Dale (MSU) for kindly providing access to their lab’s resources at the onset of this project for generation of relevant plasmid DNA and cellulose substrates. Special thanks to Leonardo Sousa for conducting ammonia pretreatment to generate cellulose III. BN would like to thank Chandra Bandi for his assistance with setting up methods crucial for execution of this project in the early phase. BN and JMY would like to thank Dr. Ashutosh Mittal (NREL) for his help with XRD measurements of Avicel cellulose derived nanocrystals. BN would also like to thank Dr. Jeff Linger at NREL for providing access to lab resources to perform QCM-D experiments. NR, CJF, and NK would like to thank Aresty Research Center at Rutgers University, for enabling research assistantships at Chundawat lab which led to their contributions to this study. BN would also like to thank Dr. Archana Jaiswal (Nanoscience instruments) for valuable discussions regarding the interpretation of QCM-D data. Conflict of interest note: SPSC declares a conflict of interest and competing financial interest(s) having filed two patent applications on pretreatment processes to produce cellulose-III enriched cellulosic biomass for biofuels production (US20130244293A1and WO2011133571A2). All other authors declare that they have no competing interests.

## Supplementary Online Material

Supporting Information (SI) appendix pdf is provided online with materials and methods/text and supplementary results (as SI Figures S1-S9, Tables T1-T3).

