## Supplementary material for "Reduced Type-A Carbohydrate-Binding Module Interactions to Cellulose Leads to Improved Endocellulase Activity": SI Appendix

**Number of SI pages in current PDF (SI-Supporting Information):** 20

**Number of SI figures in current PDF (SI-Supporting Information):** 9

**Number of SI tables in current PDF (SI-Supporting Information):** 3

### SI Appendix Materials and Methods:

***Reagents***: Reagents were procured from multiple sources as outlined here. Avicel cellulose I was obtained from Sigma Aldrich under the label Avicel PH-101. Avicel cellulose III was kindly prepared by Dr. Leonardo DaSousa (Michigan State University) using liquid ammonia pretreatement for 30 minutes at $90^0$C using 6:1 ammonia to water loading as previously described (Sousa et al., 2019). Phusion Master Mix, Dpn1 and Ni-NTA resin were obtained from ThermoFisher Scientific. Chemically competent cells were procured from various vendors: E. cloni 10g cells from Lucigen (Madison, WI), *E. coli* BL21-CodonPlus-RIPL [λDE3] from Stratagene (Santa Clara, CA) and Rosettagami 2 [DE3] from Novagen (Santa Clara, CA. pEC-GFP-CBM3a and pEC-CelE-CBM3a vectors were kindly provided by Dr. Brian Fox (University of Wisconsin, Madison)

***Synthesis and cloning of wild-type CBM genes into pEC-GFP and pEC-CelE vectors***: *E. coli* expression vectors pEC-GFP-CBM3a and pEC-CelE-CBM3a were kindly provided by the Fox lab (UW Madison) (Whitehead et al., 2017), which was used as the plasmid backbone to insert CBM1 (*Trichoderma reesei*) and CBM64 (*Spirochaeta thermophila*) genes. Plasmid maps for pEC-GFP-CBM3a and pEC-CelE-CBM3a are outlined in **SI Appendix Figure S1** and nucleotide sequence information for CBMs tested in this study are provided in the **SI Appendix Table T1.** *E. coli* codon optimized genes encoding CBM1 and CBM64, with additional flanking *AflII* and *BamHI* restriction sites, inserted into a standard pUC57-Kan vector were ordered from Genscript USA Inc (Piscataway, NJ). CBM1 and CBM64 genes were then transferred from the parent pUC57 vector to pEC-GFP and pEC-CelE vectors using polymerase incomplete primer extension (PIPE) based cloning approach as described elsewhere (Klock et al., 2009; Lim et al., 2014). Briefly, PCR amplification of destination pEC vector and CBM insert gene was conducted using Herculase II Fusion DNA polymerase and the respective PIPE primer pairs. After PCR, respective CBM PIPE reaction product aliquots (2 µL) were mixed together and immediately transformed into competent *E. coli* E. cloni 10G cells (Lucigen, Madison, WI). Vector and insert PCR reaction products were then mixed together in various ratios and immediately transformed into competent *E. coli* E. cloni 10G cells (Lucigen, Madison, WI). Presumptively positive colonies were then picked using colony screening and the plasmids extracted from these colonies were sequence verified by at the UW Biotechnology Center (or at Genscript, Piscataway, NJ). Transformed strains were stored as 20% glycerol stocks and maintained at -80 ˚C, while all relevant pEC-GFP-CBM plasmids were also maintained at -80 ˚C for long-term storage.

***Site-directed mutagenesis for generation of alanine mutants***: pEC-GFP-CBM3a/CBM64 and pEC-CelE-CBM3a/CBM64 plasmids were generated as described above and used for all mutagenesis experiments. Forward and reverse primers were designed based on QuikChange$^{TM}$ site-directed mutagenesis protocol (Braman et al., 1996), to include a codon for alanine instead of the corresponding aromatic residue. Briefly, PCR amplification was performed by mixing 10 ng plasmid DNA and 0.5 µM forward and reverse primers, with 5 µl Phusion master mix and DNase/Rnase free water to make up reaction volume to 10 µl. 1 ul Dpn1 was then added to the reaction mixture to digest native methylated wild-type plasmid DNA and then transformed into 200 µl *E.coli* E.cloni 10G cells. Colony screening and DNA sequence verification were then performed as described in the previous section.

***Protein expression and purification:*** Sequence-verified plasmid DNA was transformed into *E. coli* Rosettagami 2 [λDE3] for GFP-CBM1 and BL21-CodonPlus-RIPL [λDE3] for CelE-CBM3a/CBM64 and GFP-CBM3a/CBM64 wild-type and mutants. Transformed strains were stored as 20% glycerol stocks and maintained at -80 ˚C for long-term storage. These glycerol stocks were then used to inoculate 25 ml LB media in culture tubes, with an effective kanamycin concentration of 50 µg/ml. 25 ml overnight grown starter culture thus obtained, was then used to inoculate 500 ml LB media in 1 L shake flasks with 50 µg/ml kanamycin and grown at 37° C until exponential phase ($OD_{600}$ ~ 0.6-0.8). Protein expression was then induced using 0.5 mM IPTG at 25° C for 24 hours. Cell pellets were collected by centrifuging the liquid cultures at 10000 rpm for 15 minutes. Cell pellets were lysed using 15 ml cell lysis buffer (20 mM phosphate buffer, 500 mM NaCl, 20% (v/v) glycerol, pH 7.4), 200 ul protease inhibitor cocktail (1 uM E-64 (Sigma Aldrich E3132), 0.5 mM Benzamidine (Calbiochem 199001) and 1 mM EDTA (Fisher Scientific BP1201)) and 15 ul lysozyme (Sigma Aldrich, USA) for every 3 g cell pellet. The cell lysis mixture was sonicated using Misonix™ sonicator 3000 for 5 minutes of total process time at 4.5 output level and specified pulse settings to avoid sample overheating (pulse-on time: 10 seconds and pulse-off time: 30 seconds). The cell lysate was then centrifuged at 20000 rpm, 4 C for 45 minutes to separate cell debris from soluble cell lysate containing the protein of interest. Since all expressed proteins contained an N-terminal 8X-His tag (see **SI Appendix Figure S1**), immobilized metal affinity chromatography (IMAC) was then performed using His-Trap FF $Ni^{+2}$-NTA column (GE Healthcare) attached to BioRad™ NGC system. Briefly, the column was equilibrated in buffer A (100 mM MOPS, 500 mM NaCl, 10 mM Imidazole, pH 7.4) at 5 ml/min for 5 column volumes, followed by soluble cell lysate loading at 2 ml/min and elution using buffer B (100 mM MOPS, 500 mM NaCl, 500 mM Imidazole, pH 7.4). Electrophoretically pure protein was desalted into 10 mM MES, pH 6.5 and >90-95% purity verified using SDS-PAGE. For GFP-CBM1, a second step of purification had to be carried out using cellulose affinity chromatography as described elsewhere (Chundawat et al., 2020; Lim et al., 2014).

***GFP-CBM pull-down binding assays with cellulose allomorphs:*** Avicel cellulose-I was pre-treated at 90° C for 30 minutes to obtain cellulose-III as described previously (Sousa et al., 2019) and kindly provided by Dr. Leonardo Sousa (Courtesy of Dale Lab at MSU). The change in crystalline form was characterized using powder XRD and FTIR as discussed in previous papers from our group (Chundawat et al., 2011; Sousa et al., 2019). Binding assays were performed with at least six replicates, in 300 µl 96-well round bottomed polypropylene plates (USA Scientific). Each microwell had 200 µl total reaction mixture comprised of an appropriate volume of protein dilution to reach a certain effective concentration (0–200 µg/ml for measuring partition coefficient and 0–500 µg/ml for obtaining the full binding isotherm), 2.5 mg of Avicel cellulose-I or 10 mg of Avicel cellulose-III, an effective BSA concentration of 2.5 mg/ml to prevent non-specific protein binding while maintaining an effective buffer concentration of 10 mM MES (pH 6.5). Control microwells without cellulose were also included to obtain total protein concentration after accounting for protein loss to denaturation or non-specific binding to the microwells. The microplate was then sealed with a plate mat and shaken in an end-over-end mixing fashion in a USA Scientific hybridization oven at 5 rpm for 60 minutes at room temperature. Never-shaken control microwells were prepared similar to shaken control microwells described previously and used to obtain the calibration curve relating GFP fluorescence and known protein concentration. After 1 hour, the microplates were centrifuged at 2000 rpm for 2 minutes using an Eppendorf™ 5810R centrifuge, to separate cellulose from soluble supernatant. 100 µl soluble supernatant was picked up from each microwell using an 8-channel micropipette and transferred to opaque microplates for GFP fluorescence to be read at the following settings: 480 nm excitation, 512 nm emission with 495 nm cut-off using Molecular Devices™ UV spectrophotometer.

***Analysis of GFP-CBM pull-down binding assay data**:* After preliminary data analysis in Microsoft Excel™ to obtain free protein (μM) and bound protein concentrations (μmol/g cellulose), linear regression in Origin was used to get the partition coefficient. For full scale binding assays, the data was fit to Langmuir one-site model using the non-linear curve fitting tool in Origin. Levenberg-Marquardt algorithm with a tolerance of 1e-9 was used for curve fitting. Complex binding models such as Langmuir two-site model or Langmuir-Freundlich model were not used, to avoid overfitting errors as described in our recent study (Chundawat et al., 2020).

***CelE-CBM enzymatic hydrolysis assays with cellulose allomorphs:*** All hydrolysis assays were performed in 300 µl 96-well round bottomed polypropylene plates (USA Scientific), with at least five replicates for both reaction mixtures and blanks. Equimolar enzyme loading of 0.26 µmol enzyme per g cellulose (corresponds to 15 mg enzyme per g cellulose loading for CelE-CBM64 wild-type) was used for all enzymes to ensure consistency across mutants. Each microwell had 200 µl total reaction mixture comprised of an appropriate volume of protein dilution, 5 mg of Avicel cellulose-I or cellulose-III while maintaining an effective buffer concentration of 50 mM MES (pH 6.5). Enzyme and substrate only blanks were included to adjust for background interference from any of the reaction components. The microplates were then sealed using a plate mat and shaken end-over-end in a hybridization oven (USA Scientific) at 60° C and 5 rpm. The plates were recovered after 24 hours of incubation and reducing sugar estimation was performed using dinitrosalyclic acid (DNS) based method as described previously (Liu et al., 2020).

***Homology modeling of CBM64 structure:*** Structures for model CBMs from family 64 have been published already (**PDB codes** 5E9P and 5LU3). These constructs share ~90% and ~79% sequence similarity with the CBM64 construct used in this study. Hence, a reliable homology model of our CBM64 construct could be obtained via I-TASSER using the published structures as templates (Yang et al., 2016). Planar aromatic residues were then identified using multiple sequence alignment of our construct with the sequences of published structures.

***Reversibility testing for GFP-CBM binding to cellulose substrates:*** Binding assays to test the reversibility of CBM binding to cellulose allomorphs, were setup using a method similar to the partition coefficient binding assay reported in main manuscript with a smaller total protein concentration range (0–100 µg/ml). After the completion of partition coefficient assay and measuring GFP fluorescence of supernatant, 100 µl of reconstitution mixture (2.5 mg/ml BSA + 10 mM MES (pH 6.5)) was added to the original reaction mixtures and shaken controls. The microplate was then sealed again with a 96-well plate mat and equilibrated at 5 rpm for 60 minutes room temperature. The microplate was then centrifuged at 2000 rpm for 2 minutes using an Eppendorf™ 5810R centrifuge, to separate cellulose from soluble supernatant. 100 ul soluble supernatant was picked up from each microwell using an 8-channel micropipette and transferred to opaque microplates for GFP fluorescence to be read at the following settings: 480 nm excitation, 512 nm emission with 495 nm cut-off using Molecular Devices™ UV spectrophotometer. The new data set obtained is labeled as re-equilibration data set. If the original and re-equilibrated data sets fall roughly along the same straight line, CBM binding to cellulose would be considered reversible. A similar protocol was used in one of our previous studies (Lim et al., 2014).

***Preparation of Avicel cellulose nanocrystals through acid hydrolysis for QCM-D:*** Nanocrystals were prepared from both Avicel cellulose I and Avicel cellulose III using the same procedure. Briefly, 2 g of Avicel was added to 70 ml 4 N HCl in a glass beaker and placed in a pre-heated water bath at a temperature of 80° C. The slurry was stirred every half hour using a spatula to ensure cellulose is well suspended. After 4 hours of reaction time, the acid hydrolysis mixture was diluted with 50 ml deionized water. The slurry was then aliquoted into 50 ml centrifuge tubes, with 40 ml slurry in each tube and centrifuged at 1600xg for 10 minutes. The supernatant was decanted, and the cellulose pellet was washed with 10 ml deionized water. The wash steps were repeated, and the supernatants were discarded until they turned hazy around pH 3.3. The haziness of supernatant indicates evolution of cellulose nanocrystals and hence these supernatants were collected into a separate bottle for future usage.

***Preparation of cellulose thin films for QCM-D:*** This procedure was developed based on similar previous studies investigating cellulose hydrolysis and binding of cellulases using QCM-D (Brunecky et al., 2020). 4.95 MHz quartz crystal sensors (0.55" diameter), with $SiO_2$ coating were purchased from Filtech (product code QSX0303). The sensors were first rinsed in water, followed by ethanol and then blow-dried. The sensors were then immersed in 0.02% PDADDMAC (Sigma Aldrich 409022), which serves as an anchoring layer for cellulose, for 1 hour at 25°C with orbital mixing. This was followed by washing with deionized water for 1 hour with orbital mixing at 25°C. The sensors were then blow-dried and spin-coated using a pre-cycle spin for 3 seconds at 1500 rpm, followed by a spin cycle for 60 seconds at 3000 rpm. This spin coating step was repeated 10-20 times to obtain a uniform cellulose film thickness of ~20 nm (as measured using the QSoft software using Sauerbrey model). In a previous study (Brunecky et al., 2020), a lesser number of spin-coating steps was used to achieve the same thickness, however, this difference may be related to the nanocrystal slurry concentration and hence needs to be optimized accordingly.

***Quartz crystal microbalance with dissipation (QCM-D) based CBM-cellulose binding assay:*** Binding assays were performed using QSense E4 instrument (NanoScience Instruments). Quartz sensors with cellulose thin films were mounted and equilibrated with buffer (10 mM MES pH 6.5) at a flow rate of 100 µl/min for 10 minutes using a peristaltic pump. The flow of buffer was then stopped, and the cellulose films were left to swell overnight in buffer. The frequency and dissipation changes were tracked for all harmonics and the cellulose films were considered amenable to binding studies if the third harmonic stabilized after overnight equilibration in the buffer. Proteins of interest were diluted to a concentration of 1 µM beforehand and flown over the sensors at a flow rate of 100 µl/min for 10 minutes. All proteins tested, attained saturation within 10 minutes as noticed from frequency and dissipation traces. The CBM-cellulose system was left to equilibrate for at least 30 minutes. Unbinding of proteins was tracked by flowing 10 mM MES pH 6.5 buffer at 100 µl/min for at least 30 minutes. The sensors were finally treated with 5% contrad solution followed by deionized water at 100 µl/min for 10 minutes, to remove any traces of protein left in the tubing. The frequency and dissipation traces were analyzed using an in-house data analysis routine based on binding and unbinding equations derived as shown below.

***Analysis of QCM-D based binding assay data to obtain kinetic parameters:*** A first-principles approach was used to derive analytical equations that define the CBM binding behavior under the conditions used in QCM assay. The analysis is divided into two sections: (i) binding regime, and (ii) unbinding regime. Different assumptions hold significance under the two regimes and these assumptions are outlined in the following discussion.

$$E + C \underset{K_{off}}{\overset{K_{on}}{\rightleftharpoons}} EC$$

*Definitions and Units:* E – Free protein concentration (μM)

C – Total number of protein binding sites available on cellulose film (μmol)

EC – Number of binding sites occupied by protein (μmol)

$K_{on}$ – Association rate constant ($sec^{-1}\mu M^{-1}$)

$K_{off}$ – Dissociation rate constant ($sec^{-1}$)

N – Total number of binding sites available on deposited cellulose film in μmol (This could vary with the thickness of cellulose film being deposited)

$$\frac{d[EC]}{dt} = K_{on}[E][C] - K_{off}[EC]$$

$$\frac{d[EC]}{dt} = K_{on}[E](N - [EC]) - K_{off}[EC]$$

<u>Binding regime:</u>

*Assumption:* If we assume that the protein is in excess, then the amount of free enzyme can be assumed to be constant.

$$K_{on}[E] = K_{on}^{*}$$

$K_{on}^{*}$ – Pseudo Association rate constant ($sec^{-1}$)

Substituting $K_{on}^{*}$ in the previous equation,

$$\frac{d[EC]}{dt} = K_{on}^{*}(N - [EC]) - K_{off}[EC]$$

$$\frac{d[EC]}{dt} = K_{on}^{*}N - [EC](K_{on}^{*} + K_{off})$$

Upon integration, this gives rise to

$$\boxed{[EC] = A(1 - e^{-(K_{on}^{*} + K_{off})t})}$$

where $A = N/(1 + \frac{K_{off}}{K_{on}^{*}})$

The equation highlighted above can be used for fitting the binding regime of QCM data under the following conditions:

1. $K_{on}^{*}$ will be a true representation of $K_{on}$ only when the enzyme is in excess. Hence, at least for the wild-type protein, it will be good to test the impact of protein concentration on $K_{on}^{*}$ and make sure that we pick a concentration where the relationship between $K_{on}^{*}$ and total protein concentration is linear.
2. If $K_{on}^{*} \gg K_{off}$ then above equation can be reduced to two parameters A and $K_{on}^{*}$. Under these conditions, A is also a true representation of the number of binding sites available on cellulose, however, the dependence on cellulose concentration is not necessarily captured and hence it may not be directly correlated to the number of binding sites (μmol/g) as captured by solid state depletion binding assays.

<u>*Unbinding Regime*</u>

*Assumptions:* When unbinding begins, all the free protein is first washed out by incoming buffer. Hence the rate of adsorption is severely limited. The equation then boils down to the following, which can be used to infer the dissociation rate constant ($K_{off}$).

$$\boxed{[EC] = Ae^{-(K_{off})t}}$$

In the case of GFP-CBMs on Avicel CI and CIII, the off-rates are two orders of magnitude lower than $K_{on}^{*}$. Hence, the above assumptions hold very well in this regard. However, since $K_{on}^{*}$ is a pseudo-association rate constant which includes a free protein concentration term, true $K_{on}$ was obtained using the equilibrium dissociation constant obtained from pull-down binding assays and $K_{off}$ obtained from the analysis of QCM-D data.

### SI Appendix Figures:

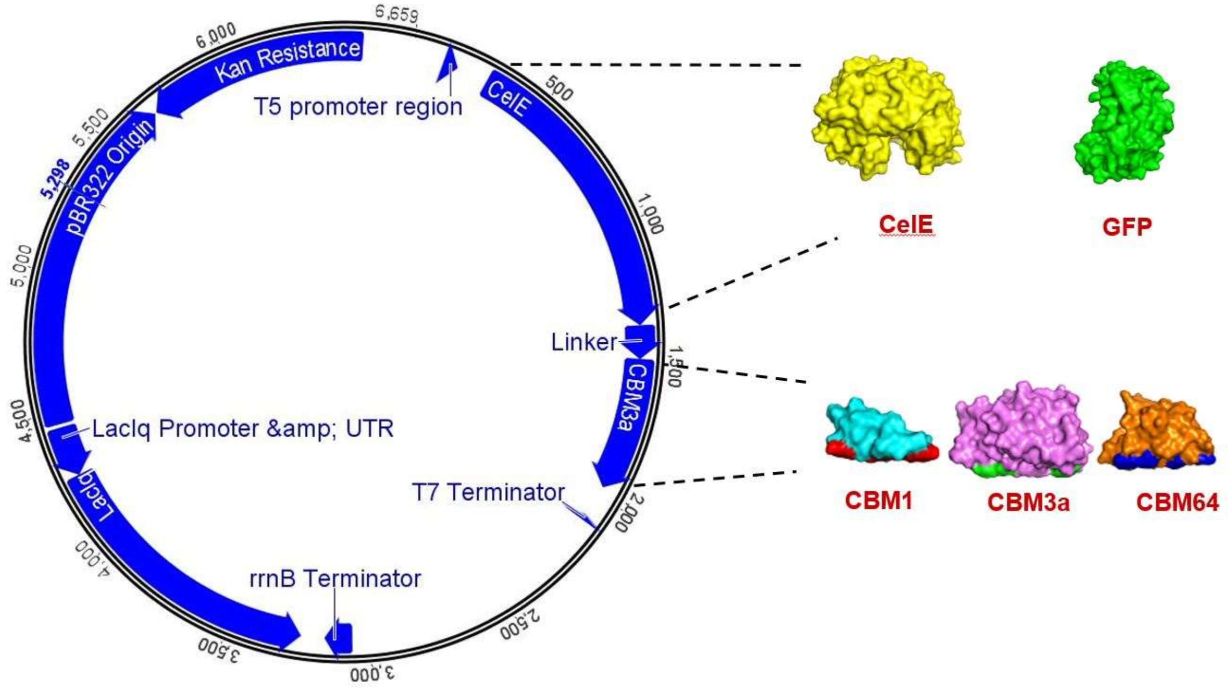


**Figure S1:** Plasmid map for pEC-CelE-CBM3a indicating the T5 promoter region followed by CelE-Linker-CBM3a and ending with T7 terminator. pEC-GFP-CBM3a vector consists of GFP gene in place of CelE, with everything else remaining unchanged.

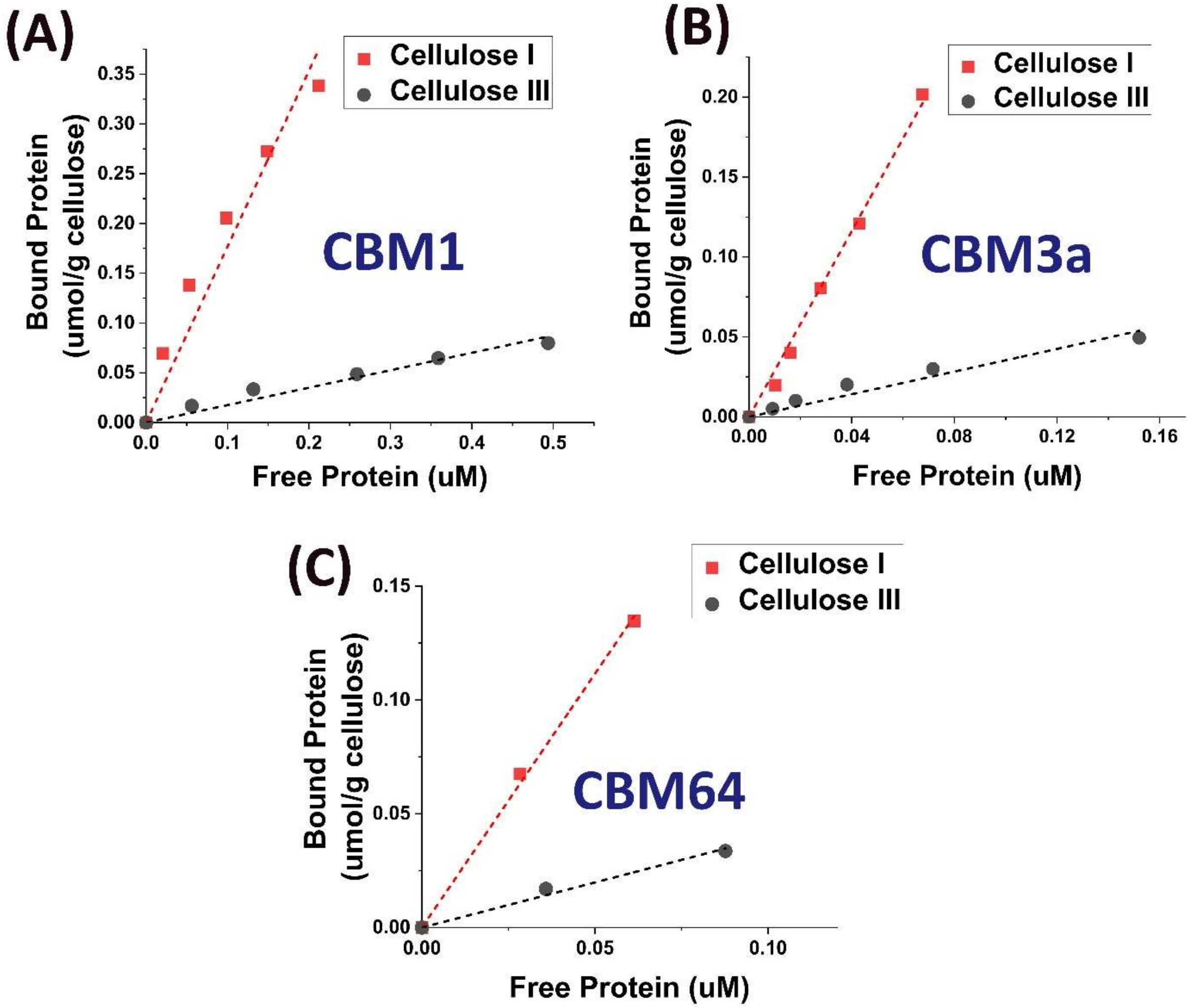


**Figure S2:** Raw data for binding of (A) GFP-CBM1 (B) GFP-CBM3a and (C) GFP-CBM64 to cellulose I (red squares) and cellulose III (black dots). Binding assays were conducted at multiple protein loadings and each data point was obtained based on at least three replicates. Red and black dotted lines represent partition coefficient (reported in **Figure 2** of main manuscript) linear fits for cellulose I and cellulose III respectively.

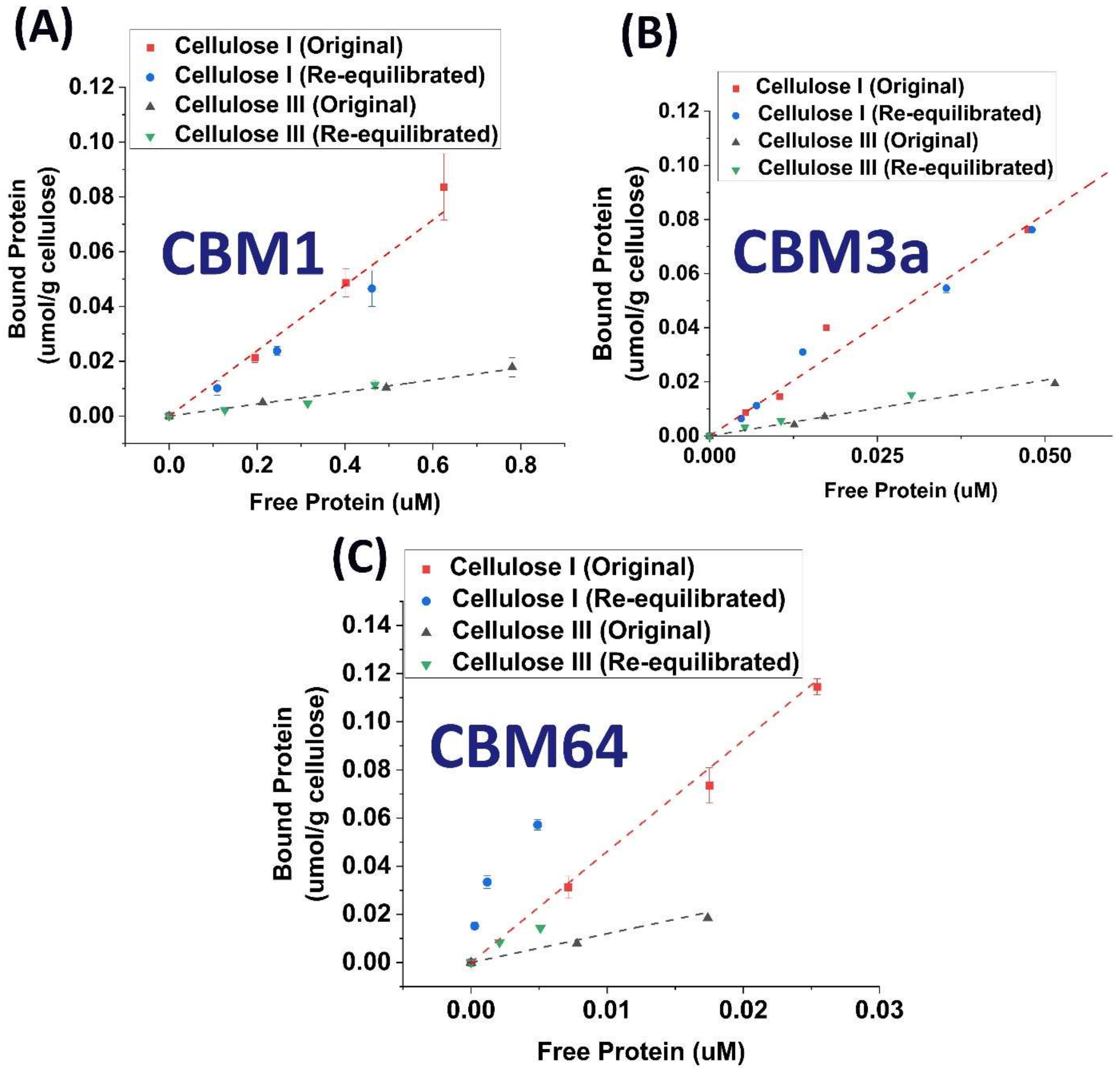


**Figure S3:** Raw data for binding assays to test reversibility of (A) CBM1, (B) CBM3a, and (C) CBM64 binding to cellulose I and cellulose III. Error bars represent standard deviations based on at least three replicates and individual data points represent equilibrium reached at a given protein loading. Red and black dotted lines represent linear fits joining original cellulose I and cellulose III data points respectively. Blue circles and green inverted triangles represent data points obtained after re-equilibration for binding to cellulose I and cellulose III respectively.

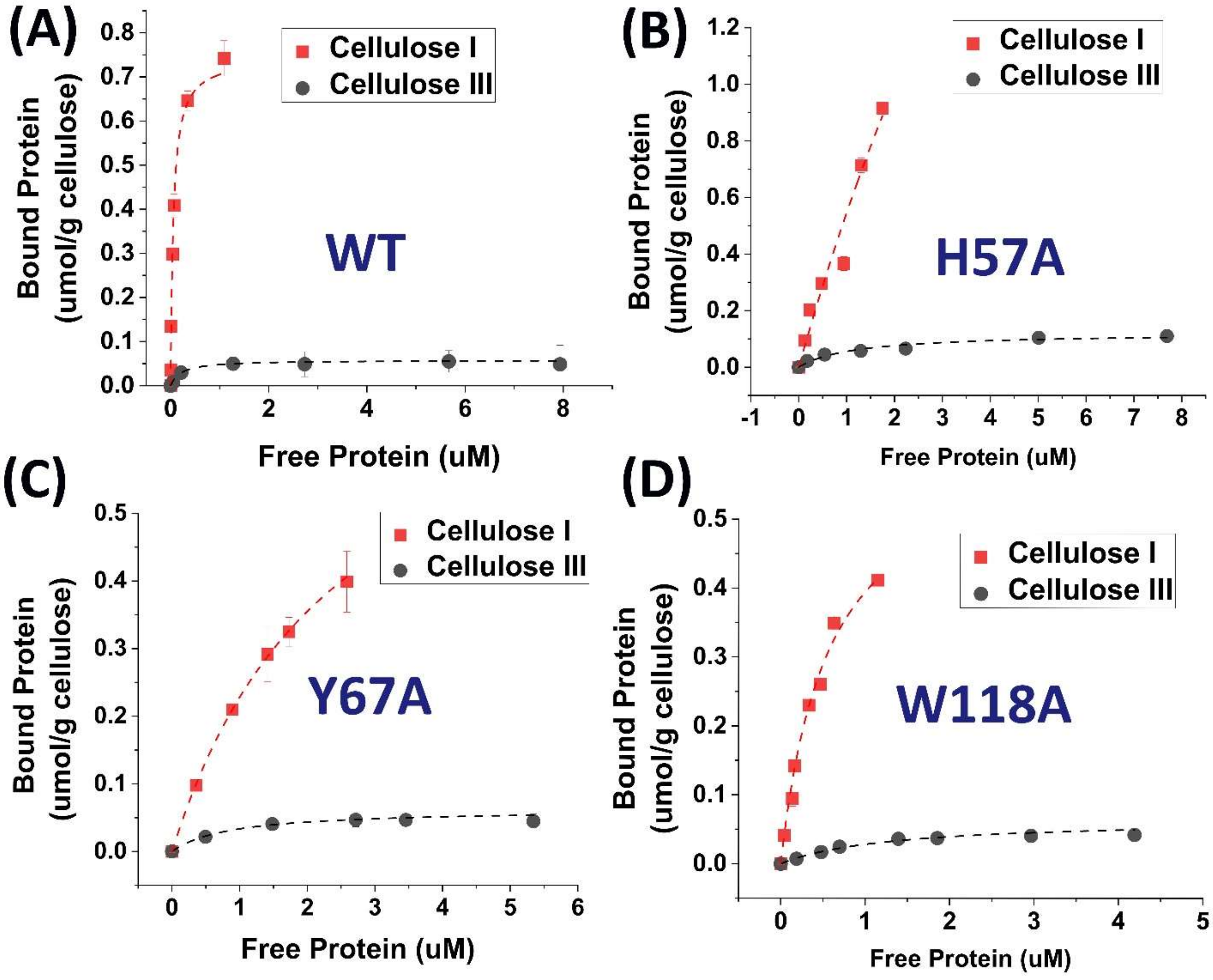


**Figure S4:** Raw data for full-scale binding assays designed to obtain total number of binding sites ($N_{max}$) and dissociation constant ($K_d$) (reported in **Table 1** of main manuscript) for binding of (A) GFP-CBM3a WT, (B) GFP-CBM3a H57A, (C) GFP-CBM3a Y67A, and (D) GFP-CBM3a W118A to cellulose I (red squares) and cellulose III (black dots). Red and black dotted curves represent Langmuir one-site binding isotherm fits for cellulose I and cellulose III, respectively. Error bars represent standard deviation from the mean based on five replicates.

Langmuir one-site binding isotherm: $[B] = \frac{N_{max}\,[F]}{K_d+[F]}$

where $[B] \rightarrow$ Bound Protein (umol/g cellulose) $[F] \rightarrow$ Free Protein (uM)

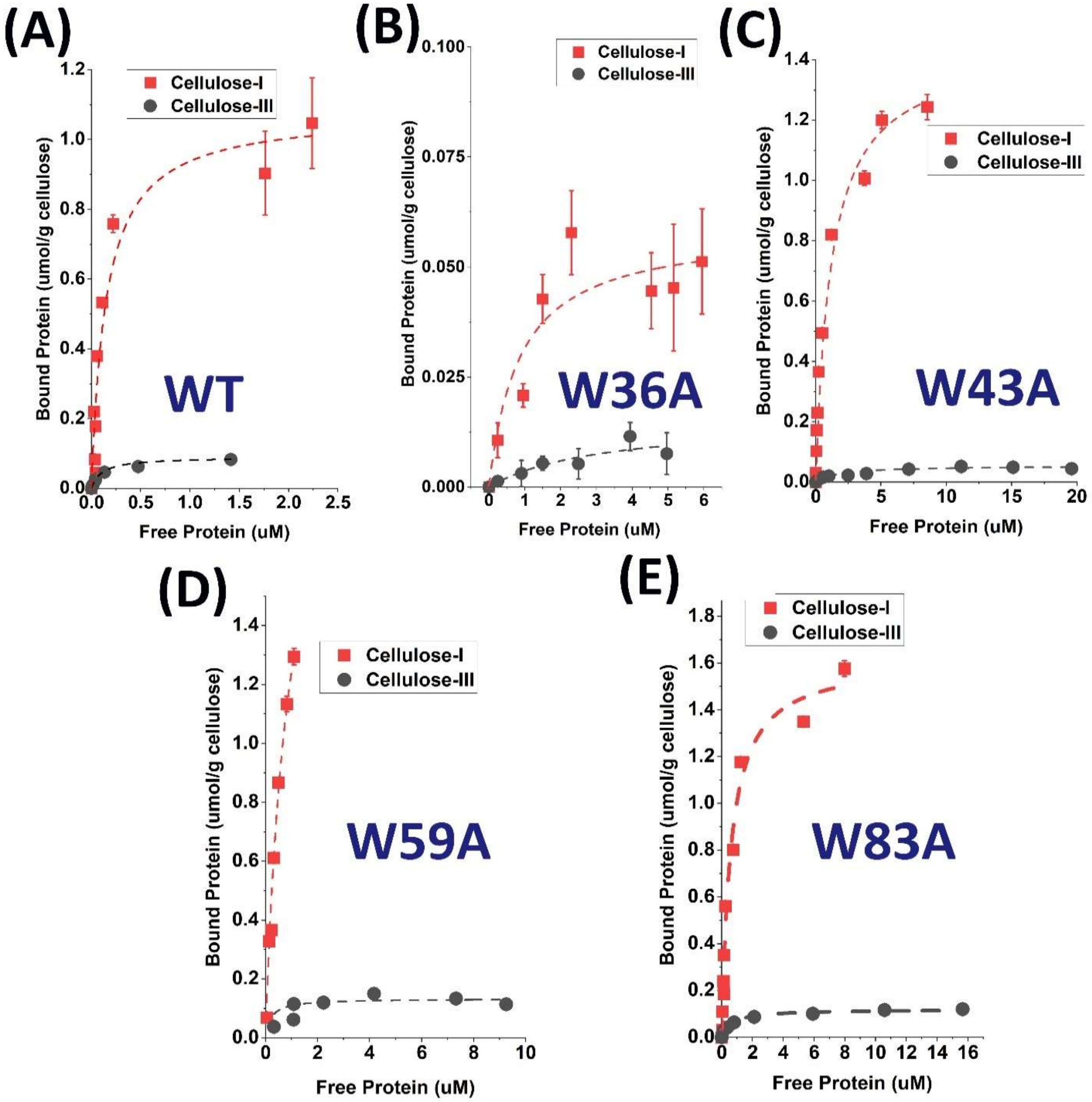


**Figure S5:** Raw data for full-scale binding assays designed to obtain total number of binding sites ($N_{max}$) and dissociation constant ($K_d$) (reported in **Table 1** of main manuscript) for binding of (A) GFP-CBM64 WT, (B) GFP-CBM64 W36A, (C) GFP-CBM64 W43A, (D) GFP-CBM64 W59A, and (E) GFP-CBM64 W83A to cellulose I (red squares) and cellulose III (black dots). Red and black dotted curves represent Langmuir one-site binding isotherm fits for cellulose I and cellulose III, respectively. Error bars represent standard deviation from the mean based on five replicates.

Langmuir one-site binding isotherm: $[B] = \frac{N_{max}\,[F]}{K_d+[F]}$

where $[B] \rightarrow$ Bound Protein (umol/g cellulose) $[F] \rightarrow$ Free Protein (uM)

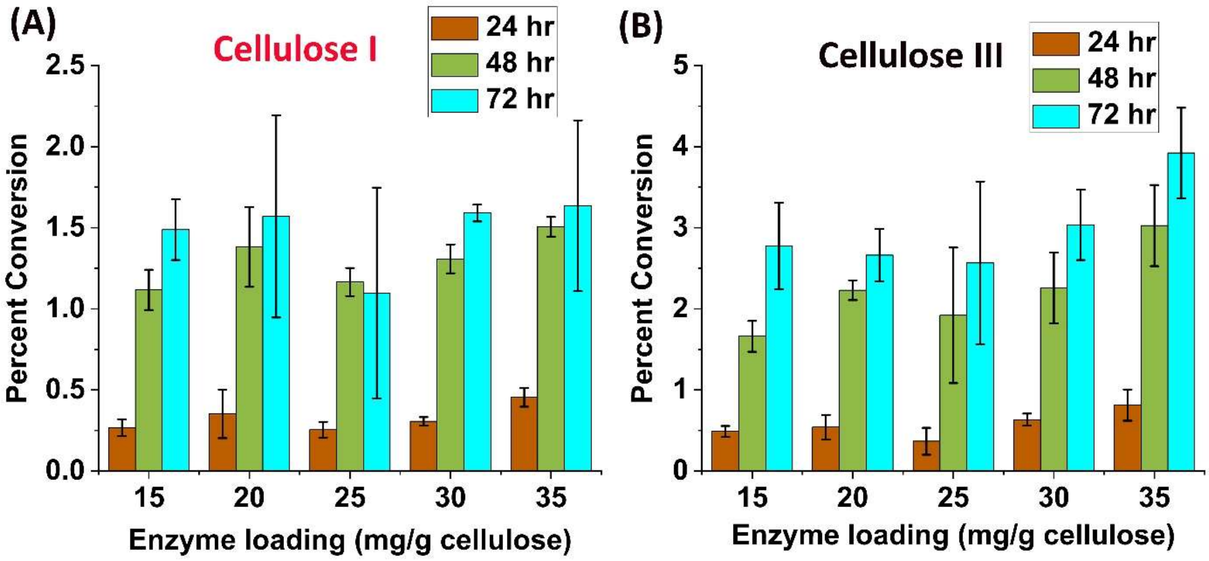


**Figure S6:** Impact of enzyme loading (15 – 35 milligram enzyme/gram cellulose) and time duration (24 – 72 hours) on the activity of CelE-CBM64-W83A towards (A) cellulose I, and (B) cellulose III. Error bars represent standard deviation from mean based on three biological replicates. Percent conversion is based on reducing sugar release measured by DNS assay and theoretical maximum conversion based on cellulose loading in reaction mixture.

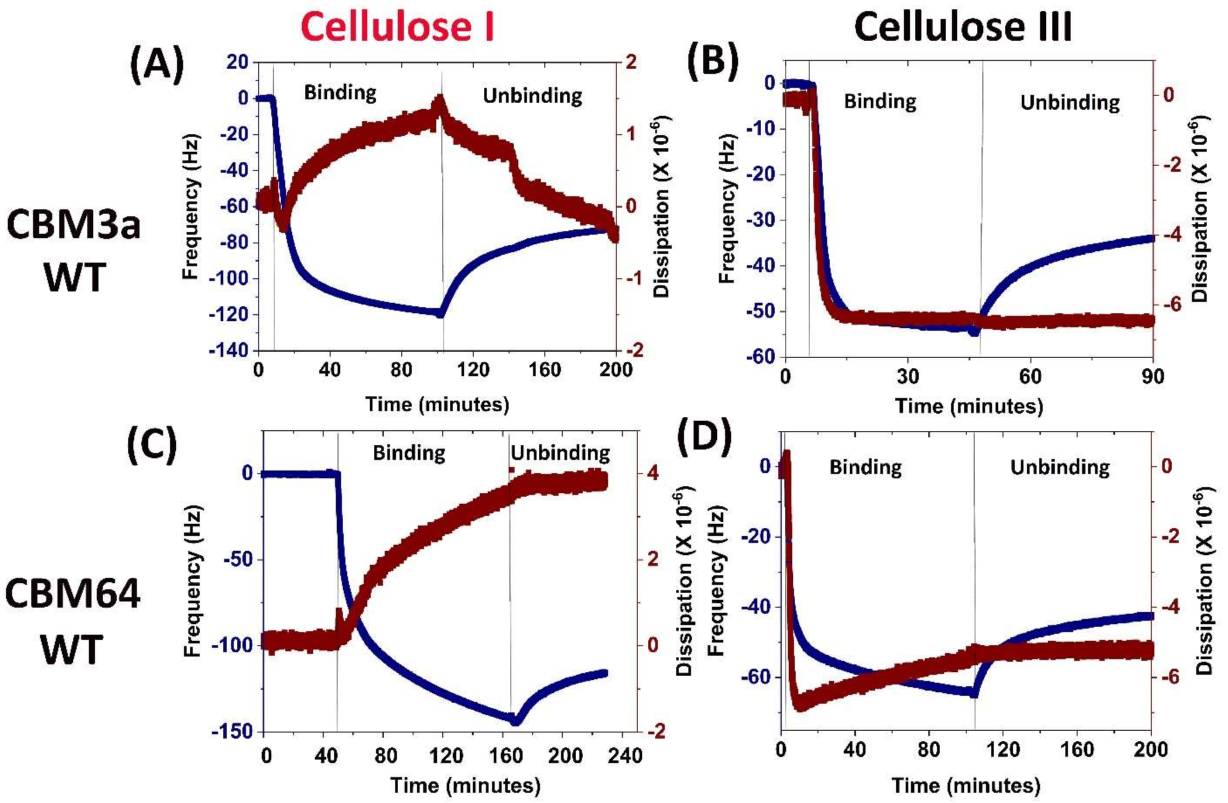


**Figure S7:** Raw data for frequency (Hz) and dissipation change vs time (minutes) as obtained from the QCM-D assay for binding of GFP-CBM3a WT on (A) cellulose I, and (B) cellulose III respectively. Similarly, the data for GFP-CBM64 WT binding to (C) cellulose I, and (D) cellulose III is reported here. The time duration of CBM binding and unbinding from cellulose surface is also indicated in each sub-figure.

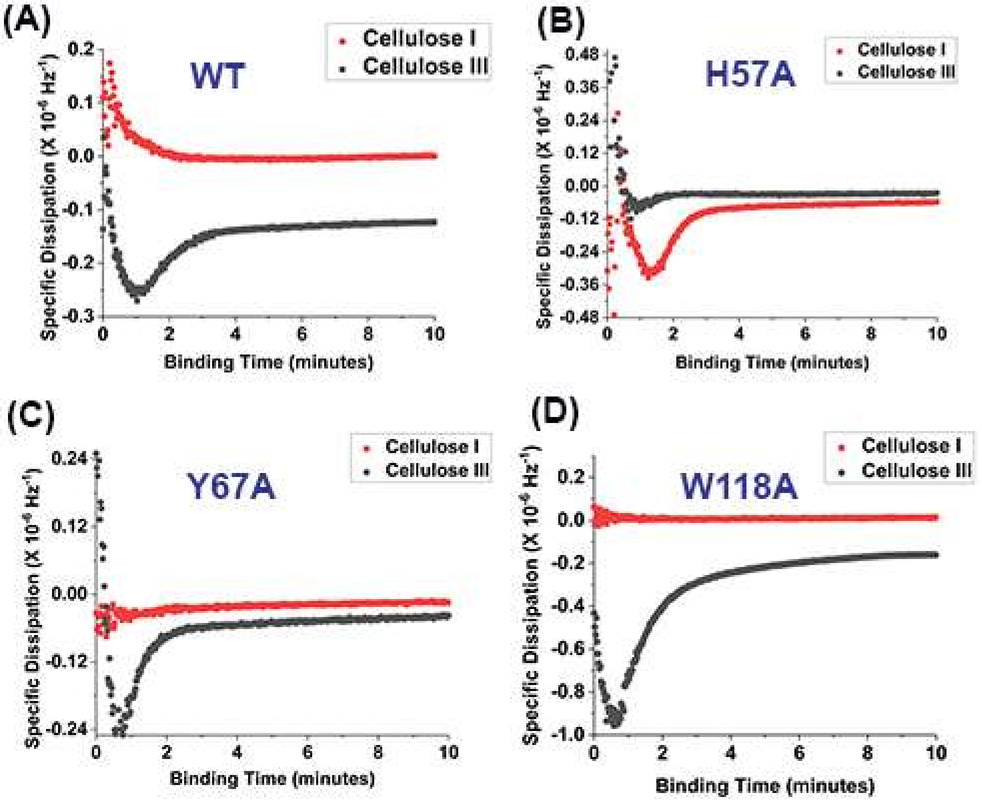


**Figure S8:** Specific dissipation for a given CBM3a mutant, (A) WT, (B) H57A, (C) Y67A, (D) W118A on cellulose I (red) and cellulose III (black). Specific dissipation is defined as the ratio of change in dissipation to negative frequency change. Sign of frequency change is flipped because reduction in frequency corresponds to an increase in adsorbed mass. One of the two data replicates used to calculate kinetic parameters, is shown here for representative purposes.

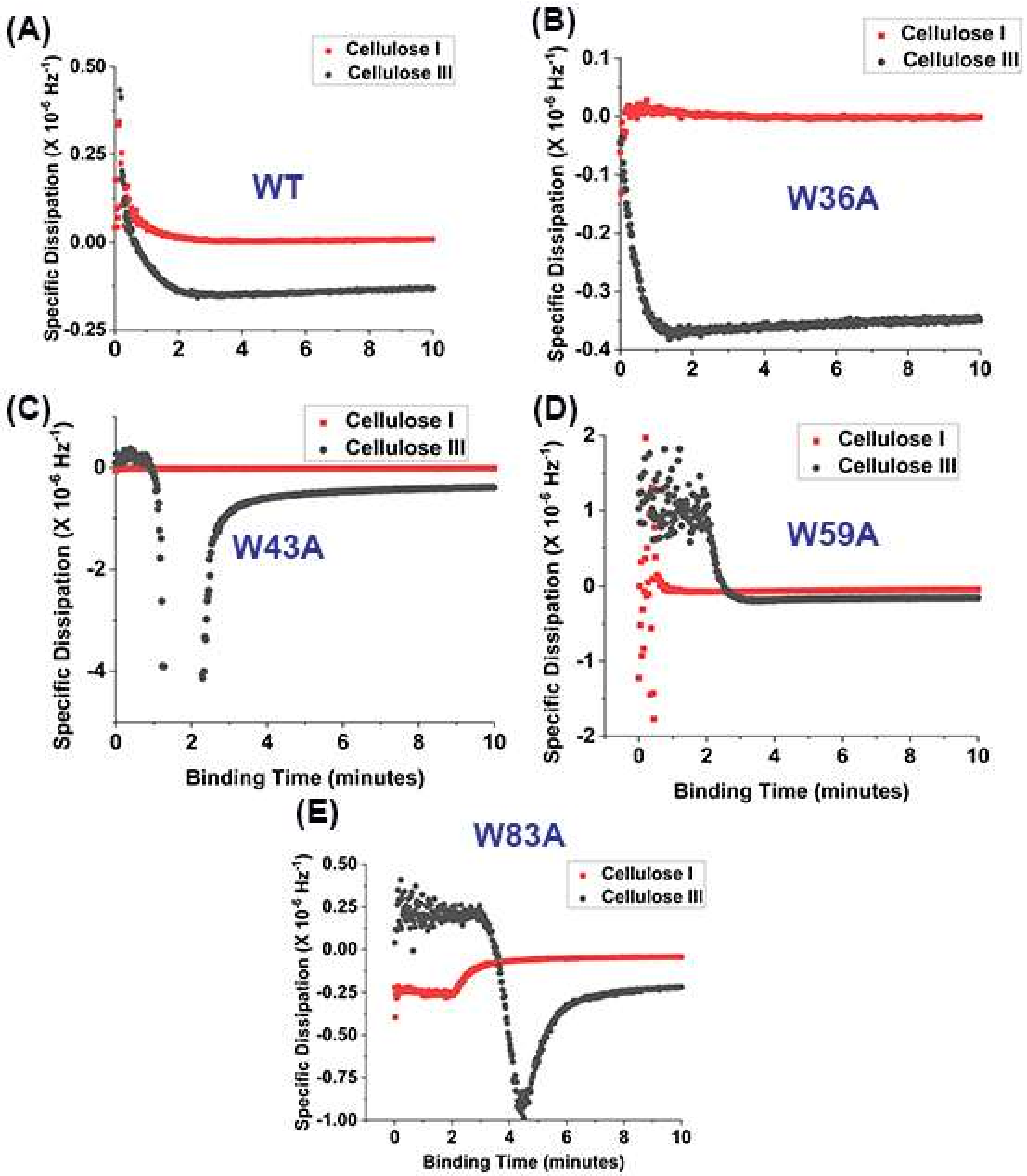


**Figure S9:** Specific dissipation for a given CBM3a mutant, (A) WT, (B) W36A, (C) W43A, (D) W59A, (E) W83A on cellulose I (red) and cellulose III (black). Specific dissipation is defined as the ratio of change in dissipation to negative frequency change. Sign of frequency change is flipped because reduction in frequency corresponds to an increase in adsorbed mass. One of the two data replicates used to calculate kinetic parameters, is shown here for representative purposes.

**SI Appendix Tables:**

| **CBM Family** | **Organism** | **Nucleotide Sequence** |
|---|---|---|
| CBM1 | *Trichoderma reesei* | CGGGTCCGACCCAGAGCCATTATGGCCAGTGCGGTGGTA<br>TTGGTTATAGCGGTCCGACCGTGTGCGCAAGCGGTACCA<br>CCTGCCAGGTGCTGAACCCGTATTATAGCCAGTGCCTG |
| CBM3a | *Clostridium thermocellum* | GTAAGCGGTAACCTGAAGGTTGAATTTTATAACTCCAAC<br>CCAAGCGACACAACGAATAGCATCAATCCGCAGTTCAA<br>AGTCACGAACACTGGCAGTTCAGCTATCGATCTGTCGAA<br>ACTGACCCTTCGTTACTACTATACGGTTGATGGCCAAAA<br>AGATCAGACCTTTTGGTGCGACCATGCAGCAATCATCGG<br>TAGCAATGGTTCTTATAACGGCATTACTTCTAATGTAAA<br>AGGCACCTTTGTGAAGATGTCAAGTAGCACCAACAATGC<br>TGATACCTACCTGGAAATTAGCTTCACGGGTGGCACACT<br>TGAACCAGGAGCCCACGTCCAGATCCAGGGCCGTTTTGC<br>GAAAAACGATTGGAGCAACTATACGCAATCAAACGATT<br>ATAGTTTCAAAAGCGCGTCTCAATTCGTAGAATGGGATC<br>AGGTGACCGCATATTTGAACGGAGTGCTGGTTTGGGGGA<br>AAGAACCAGGA |
| CBM64 | *Spirochaeta thermophila* | CCGACCCCGTCTGGCGAATATACGGCGATTGCCCTGCCG<br>TTTACCTACGATGGCGCCGGTGAATATTACTGGAAAACC<br>GACCAATTCAGCACCGATCCGAATGACTGGTCACGTTAT<br>GTCAACTCGTGGAATCTGGATCTGCTGGAAATTAACGGT<br>ACCGACTACACGAATGTGTGGGTTGCACAGCATCAAATC<br>ACGCCGGCTAGTGATGGCTACTGGTATATTCACTACAAA<br>GGCTCGTATCCGTGGTCGCATGTGGAAATCAAA |

**Table T1:** Nucleotide sequences and organism sources for wild-type CBMs from families 1, 3a, and 64 used in this study.

| Primer Name | Primer Sequence (5' --> 3') | Description |
|---|---|---|
| IP1_CBM1 | CCCGGCGAACACCCTTAAGCCGGGTCC GAC | Forward primer to amplify CBM1 gene to insert into pEC-GFP or pEC-CelE vectors |
| IP2_CBM1 | CGTTTGGGCTACTACTGCAGGTCGACT CTAGAG | Reverse primer to amplify CBM1 gene to insert into pEC-GFP or pEC-CelE vectors |
| VP1_CBM1 | GGACCCGGCTTAAGGGTGTTCGCCGG GGTATTA | Reverse primer to amplify pEC-CelE or pEC-GFP vectors for CBM1 insertion |
| VP2_CBM1 | TCTAGAGTCGACCTGCAGTAGTAGCCC AAACGAATTCGAGC | Forward primer to amplify pEC-CelE or pEC-GFP vectors for CBM1 insertion |
| IP1_CBM64 | CCGGCGAACACCCTTAAGCCGACCCC G | Forward primer to amplify CBM64 gene to insert into pEC-GFP or pEC-CelE vectors |
| IP2_CBM64 | TTCGTTTGGGCTACTATTTGATTTCCAC ATGCGACC | Reverse primer to amplify CBM64 gene to insert into pEC-GFP or pEC-CelE vectors |
| VP1_CBM64 | GTCGGCTTAAGGGTGTTCGCCGGGGTA | Reverse primer to amplify pEC-CelE or pEC-GFP vectors for CBM64 insertion |
| VP2_CBM64 | TCGCATGTGGAAATCAAATAGTAGCCC AAACGAATTCGAGC | Forward primer to amplify pEC-CelE or pEC-GFP vectors for CBM64 insertion |

**Table T2:** Primers used for SLIC based sub-cloning of CBM1 and CBM64 into pEC-GFP and pEC-CelE vectors. Insert primers were used to amplify the CBM1 and CBM64 genes synthesized by Genscript$^{TM}$ and vector primers were used to amplify pEC-GFP or pEC-CelE to eventually obtain pEC-GFP-CBM1/64 or pEC-CelE-CBM1/64 respectively.

| | **Cellulose I** | | **Cellulose III** | |
|---|---|---|---|---|
| **Mutant** | **A (x $10^{-12}$ molecules)** | **$K_{on}$* (x $10^{-2}$min$^{-1}$)** | **A (x $10^{-12}$ molecules)** | **$K_{on}$* (x $10^{-2}$min$^{-1}$)** |
| **CBM3a mutants** | | | | |
| WT | 145.6 ± 0.40 | 12.5 ± 2.1 | 97.71 ± 1.62 | 14.0 ± 1.4 |
| H57A | 64.22 ± 0.23 | 25.5 ± 6.3 | 79.80 ± 16.7 | 8.00 + 1.4 |
| Y67A | 60.88 ± 5.9 | 18.0 ± 1.4 | 49.15 ± 0.353 | 8.15 ± 0.78 |
| W118A | 106.9 ± 0.57 | 0.055 ± 0.0070 | 64.52 ± 8.37 | 14.5 + 0.021 |
| **CBM64 mutants** | | | | |
| WT | 203.7 ± 2.62 | 40.0 ± 0.0 | 91.52 ± 5.90 | 29.0 ± 1.4 |
| W36A | 75.15 ± 7.69 | 15.0 ± 4.2 | 66.70 ± 16.8 | 11.4 ± 2.4 |
| W43A | 89.20 ± 0.53 | 13.5 ± 0.7 | 57.90 ± 33.8 | 13.5 ± 2.1 |
| W59A | 87.45 ± 6.08 | 12.0 ± 0.0 | 80.48 ± 11.9 | 20.5 ± 15 |
| W83A | 117.1 ± 5.46 | 11.5 ± 2.1 | 45.86 ± 4.39 | 13.9 ± 0.070 |

**Table T3:** Binding parameters ($A, K_{on}^{*}$) from fitting of QCM-D assay data. Relationship between the number of protein molecules and binding parameters is described in terms of the following equation:

$$[EC] = A(1 - e^{-(K_{on}^{*})t})$$

where $[EC] = Number\ of\ molecules\ t = Time\ (minutes)$
